# Genomic structure, introgression, niche overlap, and morphological variation challenges species delimitation in Thymus sect. Mastichina

**DOI:** 10.64898/2025.12.16.694552

**Authors:** Diego Nieto-Lugilde, José C. del Valle, Esther María Martín-Carretié, David Doblas, Billy Williams-Marland, M. Ángeles Ortiz, Francisco Javier Jiménez-López, Javier López-Tirado, Francisco José García-Cárdenas, Regina Berjano

## Abstract

Species delimitation remains a major challenge in understanding biodiversity, particularly in recently diverged lineages, where extensive morphological variability complicates taxonomic inference. In this study, we investigate patterns of morphological and ecological variation within *Thymus* sect. *Mastichina*, a Mediterranean lineage that includes the endangered species *T. albicans*. Previous genomic analyses revealed clear differentiation between two major groups corresponding to ploidy levels, as well as additional genetic structure among diploid lineages. Here, we integrate this genomic framework with detailed analyses of floral morphology and ecological niches to assess the correspondence between phenotypic, ecological, and genetic variation. Our results reveal substantial overlap among genetic lineages at both morphological and ecological levels. Although some differentiation is associated with ploidy level, particularly in floral traits, extensive variability within several lineages obscures clear taxonomic boundaries based on discrete morphological characters. By contrast, some diploid lineages (notably the Cádiz and Algarve groups) show lower morphological variability but still both overlap in morphology and ecological niche. We morphologically characterized a recently discovered diploid lineage (Hercynian) that appears to be cryptic with the tetraploid *T. mastichina*. Patterns of admixture involving this Hercynian lineage, especially with the diploid Doñana group, contribute to pronounced phenotypic overlap, whereas no significant effect of tetraploid introgression on diploid morphology was detected. Ecological niche analyses indicate that niche differentiation is primarily associated with ploidy level rather than among diploid lineages, except for the Hercynian group. Overall, our results demonstrate a marked discordance between phenotypic variation and genetic structure, supporting the interpretation of *Thymus* sect. *Mastichina* as a complex of cryptic evolutionary lineages. These findings highlight the limitations of morphology-based taxonomy and emphasize the need for an integrative framework incorporating genome size, genomic information, and geographic distribution for species delimitation.

## Introduction

Recent advances in genetic analysis have revealed an increasing number of taxa that challenge traditional species boundaries based only on morphological criteria (Hending 2025). This may have important implications for biodiversity assessment, particularly in systems in which cryptic species, morphologically similar taxa distinguishable only at the genetic level, might remain undetected. Such hidden diversity can lead to a misestimation of species richness and undermine the design and implementation of effective conservation strategies, especially for endemic or protected taxa (Bickford et al. 2007; Poulin and Pérez-Ponce de León 2017; Struck et al. 2018). Therefore, it is essential to determine whether cryptic lineages also exhibit meaningful differences in morphological traits, ecological niches, or geographical distribution that are critical for guiding field identification and conservation priorities (Bickford et al. 2007; Hending 2025).

The Mediterranean Basin stands out as a region where the question is especially relevant, due to its unique combination of exceptional plant diversity, high levels of endemism, and increasing ecological threats (Cuttelod et al. 2008). Despite covering a small fraction of the Earth’s surface, the Mediterranean Basin hosts a remarkable diversity of habitats and supports an exceptionally high plant species richness, accounting for approximately 10% of global vascular plant diversity (Greuter 1991; Myers et al. 2000; Thompson 2005; Blondel et al. 2010; Nieto Feliner et al. 2023). Notably, around 60% of plant species in this region are endemic, underscoring the critical role of endemism in shaping its floristic diversity and originality (Greuter 1991; Melendo et al. 2003; Thompson et al. 2005). This extraordinary botanical wealth is the result of complex abiotic influences, including climatic seasonal and spatial variability, soil heterogeneity, and topographic variation, alongside a dynamic geoclimatic history shaped by tectonic activity, glaciations, and shifts in sea levels (Holzapfel et al. 1992; Matesanz and Valladares 2014; Nieto Feliner et al. 2023). In addition to these natural influences, anthropogenic pressures have also significantly impacted Mediterranean plant diversity (Médail and Diadema 2006; Thompson 2020; Peñuelas and Sardans 2021). In this context, plant conservation emerges as a critical priority for maintaining ecosystem stability, yet it faces major challenges, including limited assessments of species conservation status, insufficient funding, and gaps in knowledge related to the species’ distributions, ecology, and/or reproductive biology (Havens et al. 2014; Heywood 2017).

The Lamiaceae family is a good example of plants adapted to the summer aridity that characterize the Mediterranean climate (Matesanz and Valladares 2014; Aedo et al. 2017), with thymes (*Thymus* spp.) standing out for its remarkable diversification (Morales 1986; Melendo et al. 2003). However, this is a taxonomically complex group, since phenotypic plasticity, recent divergence, and hybridization obscures diagnostic traits, blurring the lines between closely related taxa (Morales 2002).

The genus *Thymus* L., which includes mostly circum-Mediterranean species, is taxonomically divided into eight sections (Jalas 1971; Morales 1986). Among these, the section *Mastichina* Mill. comprises two gynodioecious, aromatic species endemic to the Iberian Peninsula: *T. mastichina* L. and *T. albicans* Hoffmanns. & Link (Morales 2010). While *T. mastichina* is widely distributed across most of the Iberian Peninsula, *T. albicans* has a much more restricted geographic range, being mainly confined to the southwestern part of the Iberian Peninsula, mainly in Cádiz (Spain) and Algarve (Portugal). Indeed, *T. albicans* is listed as Critically Endangered (CR) in the Red List of Vascular Flora of Andalusia (Decreto 23/2012) and in the Red List of Spanish Vascular Flora (Moreno 2008), and as Vulnerable (VU) in the IUCN Red List (Buira et al. 2017). Taxonomic classifications have recognized two subspecies within *T. mastichina*: *T. mastichina* subsp. *mastichina* L., a tetraploid taxon widely distributed throughout most of the Iberian Peninsula, and *T. mastichina* subsp. *donyanae* R. Morales, a diploid taxon inhabiting mainly on the sandy soils of Doñana National Park (Huelva, Spain). However, recent phylogenomic studies have revealed that *Thymus* sect. *Mastichina* is more accurately divided into two well-supported clusters: one comprising tetraploid individuals, mostly with distribution corresponding to *T. mastichina* subsp. *mastichina* sensu Morales (2010), and the other consisting of diploid individuals, taxonomically classified in Morales (2010) as *T. albicans* and *T. mastichina* subsp. *donyanae*, but also some populations identified as *T. mastichina* subsp. *mastichina* (García-Cárdenas et al. 2025). According to this study, the diploid cluster can be further divided into four genetic groups, each exhibiting a clear geographic pattern: Algarve (Algarve; Portugal), Doñana (Huelva; Spain), Cádiz (Cádiz; Spain), and Hercynian (Cordoba, Spain; and Serra da Estrela, Portugal) (García-Cárdenas *et al*., 2025). Moreover, in most populations, signs of genetic introgression (i.e., the incorporation of genetic material from one species or population into another) have been detected, at both intraploidy (among diploid genetic groups) and interploidy level (between diploids and tetraploids; García-Cárdenas *et al*., 2025).

By introducing new alleles and genetic combinations, introgression may increase phenotypic variation, altering characteristics such as size, shape, structure, or coloration of vegetative and reproductive organs (Aguillon et al. 2022). This enhanced variability can lead to the emergence of novel traits or modifications of existing ones, potentially affecting ecological adaptation and population dynamics (Suarez-Gonzalez et al. 2018; Villa-Machío et al. 2024). On the contrary, introgression can also blur or modify species-specific morphological features, complicating taxonomic identification and impacting local evolutionary trajectories (Thórsson et al. 2001).

Given the cryptic genetic diversity, its mismatch with previous taxonomical works (Morales 2010), and the prevalence of introgression in the *Thymus* sect. *Mastichina* (García-Cárdenas et al. 2025), it seems necessary to study the morphological and ecological features of the group. Here, we aim to gather and analyse morphological and ecological data in the *Thymus* sect. *Mastichina* to discern patterns among ploidy levels and/or genetic clusters. More specifically, we aim to discern whether there are recognizable morphological or ecological differences between the different lineages (ploidy levels and genetic groups) that could contribute to clarify the taxonomy of the section and what is the role of introgressions in its phenotypical traits. We hypothesize that 1) the tetraploid individuals display a wider phenotypical and ecological variability than the diploid individuals; 2) the different genetic groups might display some different but overlapping morphologies and/or ecologies, and 3) introgression of the tetraploid genome into diploid populations may affect morphological, and ecological features in diploids, causing overlap in species’ features.

## Materials and Methods

To test our hypothesis, we studied multiple populations of *Thymus* sect. *Mastichina* across the Iberian Peninsula. For each population, we characterized their main morphological traits (namely floral traits commonly used in the taxonomy of the genus), their abiotic environment through soil and climate conditions, and their biotic environment through vegetation quadrats. Finally, we complemented our dataset by gathering existing information on the ploidy level and genetic group assignment of each population. These morphological and environmental datasets were evaluated using a suite of univariate and multivariate statistical approaches to test for differentiation among ploidy levels and genetic clusters. Finally, we also examined the influence of ancestry introgression on floral trait variation.

### Plant material

We collected samples from 39 populations spanning the Iberian Peninsula, encompassing the distribution of *Thymus* sect. *Mastichina*. Given that the distribution area of *T. albicans* is restricted to the southwestern region of the Iberian Peninsula, sampling effort was concentrated in this area (**Figure 1 and Table S1**). From May to June 2023, during peak flowering, we collected samples from 15 hermaphroditic and 15 female plants per population. Individuals were randomly selected in the population at approximately the same phenological stage. Marked individuals were at a minimum distance of 5 m between them to reduce the probability of sampling siblings. At least one flowering stem per individual was collected and preserved as a herbarium specimen. One sample per population was deposited in the SEV herbarium of the University of Seville (**Table S1**).

**Figure 1.**
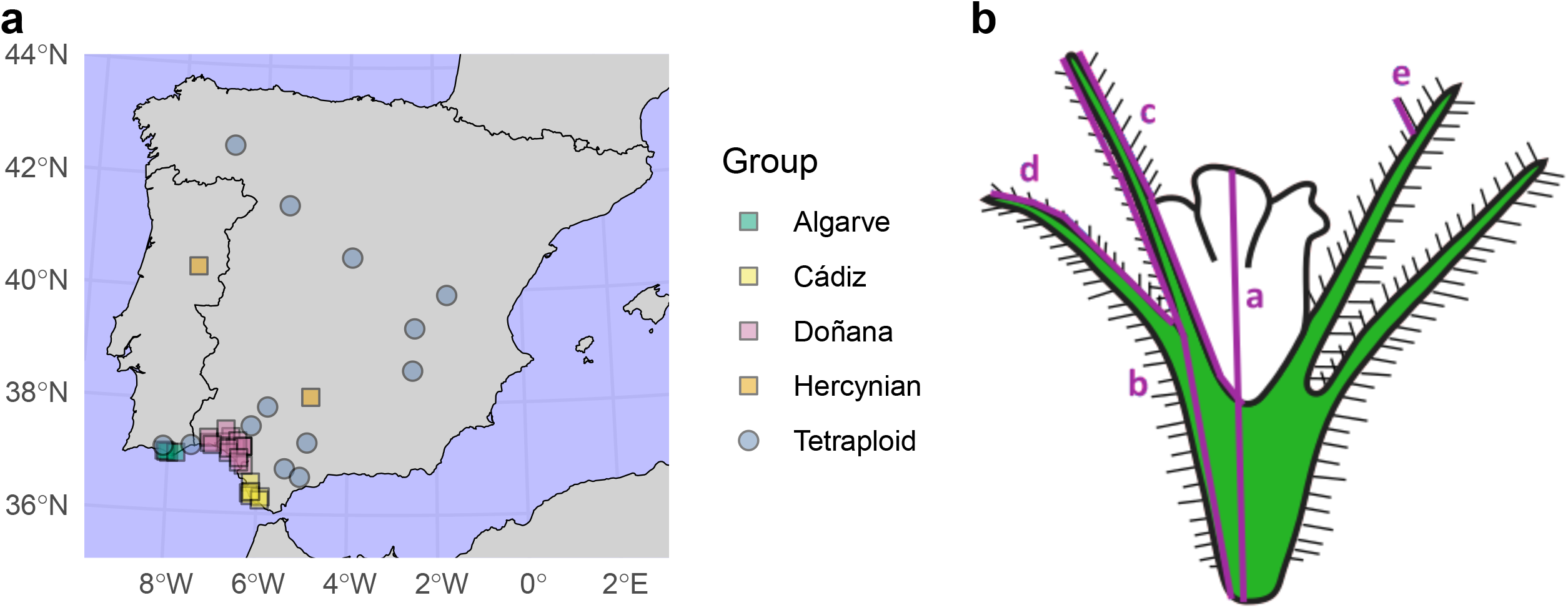
Geographical sampling of populations from Thymus sect. Mastichina and schematic representation of measured floral traits. The map (a) depict the Iberian Peninsula showing sampling locations coloured by genetic group (García-Cárdenas et al., 2025): Algarve (green), Cádiz (Cádiz), Doñana (Yellow), Hercynian and Tetraploid. The diagram represent a flower indicating the morphological traits measured: (a) corolla length, (b) calyx length, (c) length of the longest calyx tooth, (d) *length of the shortest calyx tooth*, and (e) length of the longest calyx hair. The studied populations are color-coded according to the five genetic groups identified by García-Cárdenas et al. (2025): Tetraploid (blue), Hercynian (orange), Algarve (green), Doñana (yellow), and Cádiz (pink). Circles and squares represent diploid and tetraploid locations, respectively. The map is in UTM projection for the zone 30N and datum WGS84.

### Genomic data and ploidy level

The sampled populations were the same as those used in the genomic analysis described in García-Cárdenas *et al*. (2025); consequently, we used the information from that study to obtain variables that characterize the studied individuals. Specifically, García-Cárdenas et al. (2025) estimated 2C DNA content of the same individuals of this study by flow cytometry, allowing the classification of individuals into diploids and tetraploids. Additionally, genetic clustering and ancestry estimates derived from genome-wide SNP data generated through HybSeq further separated the Tetraploid group and four diploid genetic groups: (1) Algarve, (2) Doñana, (3) Cádiz, and (4) Hercynian. Lastly, we obtained the percentage of genetic ancestry for all individuals also from ADMIXTURE analysis. With all this information we created two categorical variables (i.e., ploidy level and genetic group) and one continuous variable (i.e., percentage of genetic ancestry) for each plant individual.

### Morphological analysis

For the morphological characterization of the studied populations, we measured the following floral traits: calyx length (ca), corolla length (co), lengths of the longest calyx tooth (ltooth), length of the longest hair on calyx teeth (hair), number of flowers in the inflorescence (fl_infl), and diameter of the inflorescence (d_infl; **Figure 1 and Table S2**). Floral traits were measured in 1-7 open flowers from 3-12 collected individuals in each population (including both hermaphroditic and female individuals), yielding a total of 965 flowers. These flowers were extracted from their inflorescences and digitized with an Epson Expression 11000XL high resolution scanner. The resulting images were analysed using the software ImageJ v.1.53t (Schneider et al. 2012).

Additionally, we started a tentative analysis of microscopic features (i.e., pollen and stomata sizes). More specifically, we measured the minor and major pollen axis (d_min and d_max, respectively) and the length and width of the stomata (stoma_l and stoma_w, respectively). We measured 10 pollen grains and 10 stomata from 1-2 individuals in 11 different populations (7 populations for each feature; n = 100 and 210 for pollen and stomata size, respectively) following the preparation protocol described in Scarpeci et al. (2017) and using a Leica DFC-490 microscope. However, the preliminary results suggest that there is no different signal in the microscopic features and, hence, we did not continue fully measuring the rest of populations (**Figure S1 and Tables S2 and S3**).

### Environmental characterization of natural populations

To characterize abiotic environmental conditions in each population, we gathered climatic data and soil samples. Climate was characterized by the extended bioclimatic (Bioclim+) variables from the Chelsa Climate Database v.2.1 (Brun et al. 2022). We extracted values for aridity index (ai), the bioclim variables (bio1-19), mean climate moisture index (cmi_mean), growing degree days heat sum above 5°C (gdd5), and mean near-surface wind speed (sfcWind_mean) using the geographic coordinates of the populations recorded in the field when collecting plant samples. To reduce the multidimensionality, we performed a Principal Component Analysis (PCA) on the whole set of variables and retained the first five axis for further analysis.

We obtained edaphic variables for each studied population from a prior analysis of three soil samples from the same localities, as published in del Valle et al. (2025). Edaphic variables include pH, electrical conductivity (EC), water retention potential (WRP), water retention capacity (WRC) and organic matter (OM).

To characterize the plant community, at each population we set three quadrats of four squared meters following Lavergne et al. (2004). Vascular plants were inventoried by means of point contact for 100 points for each quadrat (20 × 20 cm grid). Coverage plant species was assessed by summing the contact points and then dividing by 100 (total number of contact points). Additionally, we completed the list of companion species by randomly walking around the quadrats (15-20 m diameters) and identifying other species absent in the quadrats. Because in some sites, the companion species were not represented in the quadrats but might be important in the community (e.g. tree or shrub species), they were also included in the inventories with the lowest coverage (i.e., 1 %).

### Statistical analyses

To explore overall patterns of variation in morphological traits and environmental variables (both bioclimatic and soil), we assessed differences between ploidy levels (diploids vs. tetraploids) and among the five genetic clusters identified within *Thymus* sect. *Mastichina*. We also explored morphological differences between sexes (hermaphrodites and female individuals). As the data did not meet normality assumptions, we used non-parametric tests (Mann-Whitney *U* for comparison between ploidy levels and Kruskal–Wallis tests, followed by Dunn’s post hoc test, for comparison between genetic groups). Additionally, to evaluate potential differences between lineages in multivariate spaces, for both morphological and environmental variables, we performed a PCA using functions from base R and retained the first two axis. Then, we visualized the PCA results using the *factoextra* v.1.0.7 R package (Kassambara and Mundt 2020). Finally, we analysed variation in calyx size (one of the main features used in taxonomy of the section) using Bayesian hierarchical linear models fitted using the *brms* v.2.23.0 R package, partitioning phenotypic variance among genetic groups, populations, individuals and residual error. Models were fitted using Hamiltonian Monte Carlo sampling, and posterior distributions of variance components and their proportional contributions to total variance were summarised using posterior means and 95% credible intervals. Separate hierarchical models were fitted for each genetic group to characterise and compare the internal distribution of variance among populations, individuals and residual components within groups.

To study differences in the ecological communities, we used Principal Coordinates Analysis (PCoA), by standardizing the species cover data with the Hellinger transformation and calculating Bray-Curtis distance between quadrats, as applied in the decostand and vegist functions in the *vegan v*.*2*.*7*.*2* R package (Oksanen et al. 2018).

Finally, to test how ancestry introgression among genetic groups affects floral morphology, we used the genomic data generated by García-Cárdenas *et al*. (2025). More specifically, we fitted regression models, using the percentage of ancestry to different genetic groups as predictors and each floral trait as a response variable. Within each of the five main genetic groups, we modelled the effect of ancestry from the remaining genetic groups, which we interpret as ancestry introgression, on trait variation. We first fitted Gaussian linear mixed models with a random intercept for sex (i.e., females and hermaphrodites) with the package glmmTMB, and compared them to models without the random effect using the corrected Akaike Information Criterion (AICc); when the random effect did not improve model fit (ΔAICc < 2), we retained the simpler linear models. In addition to multivariate models including several ancestry components simultaneously, we also fitted univariate models with a single percentage of genetic ancestry as predictor, to assess the effect of each ancestry component independently. Model assumptions were checked by visual inspection of residuals Q-Q plot.

## Results

### Floral morphology

When comparing diploid and tetraploid individuals, we found that the tetraploids exhibited significantly larger floral traits than diploids across all measured parameters (**Figure 2; Tables S2 and S3**), except for microscopic features (pollen grain and stomata size; **Figure S1**). Despite these significant differences, we discovered substantial variability and overlapping between the two ploidy levels (**Figure 2**). These results were corroborated by the overlapping position of the points’ clouds in the PCA analysis (**Figure S2**), which did not clearly separate some tetraploid and diploid individuals. The first two components of the PCA accounted for 69.2% and 12.4% of the total variation, respectively.

**Figure 2.**
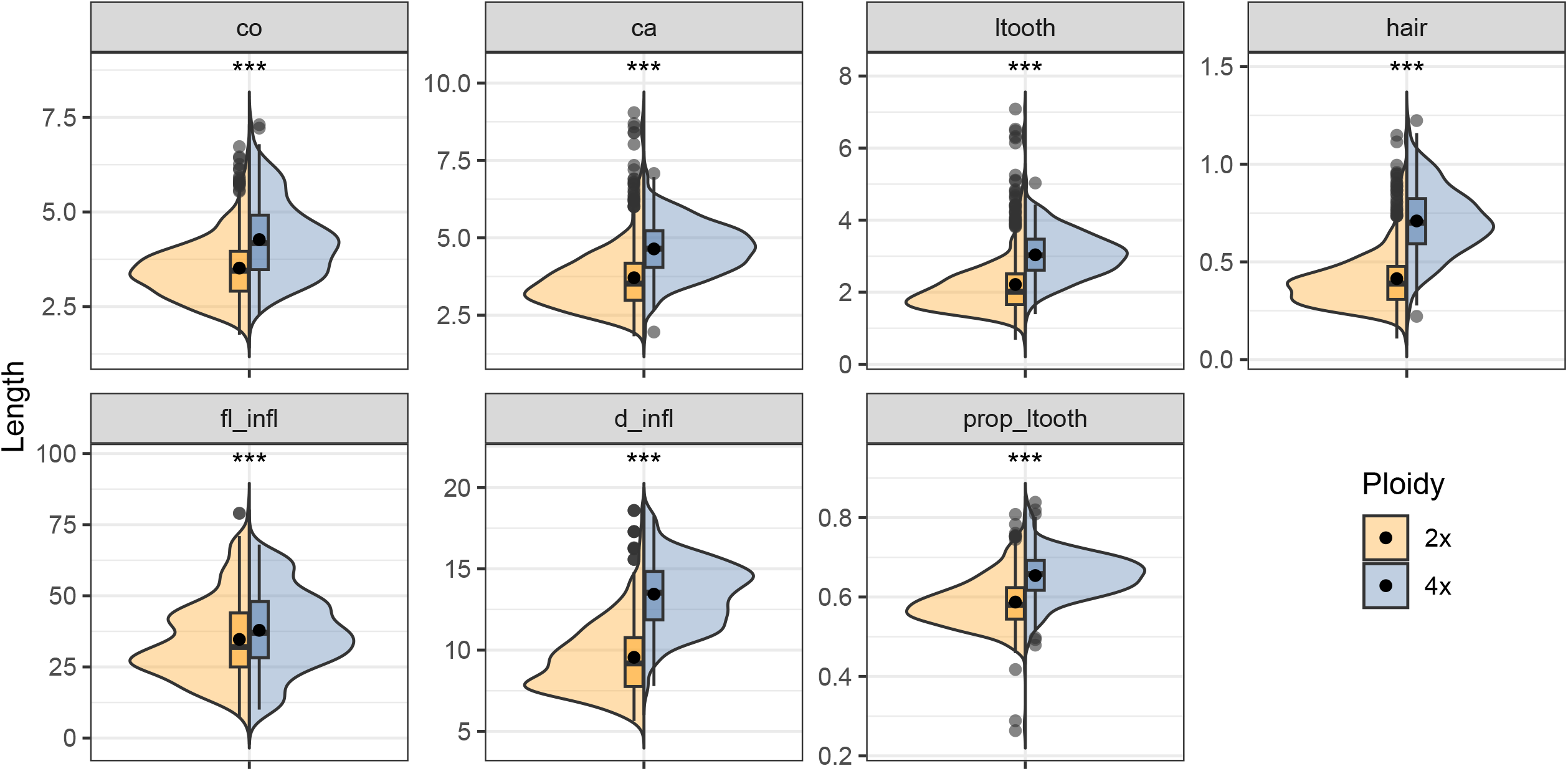
Comparison of morphological traits between ploidy levels. Violin plots combined with boxplots show the full distribution of values obtained for each measured morphological trait: corolla length (co), calyx length (ca), length of the longest calyx tooth (lteeth), length of the longest calyx hair (hair), number of flowers in the inflorescence (fl_infl), inflorescence diameter (d_infl), proportion of the calyx tooth (prop_ltooth). All metrics are in millimeters. Boxplots represent the interquartile range (IQR); the thin black line indicates 1.5× the IQR, the central line shows the median, and the black dot represents the mean sample value; black dots outside the IQR represent the outlayer values. Asterisks denote significant differences between ploidy levels (ns, non-significant; *, P < 0.05; **, *P* < 0.01; and ***, *P* < 0.001).

In general, 41% of the observed variance for calix length was attributed to differences among the five genetic groups, while 40% was attributed to differences among populations and only 14% within individuals (**Table 1**). All individuals from Algarve and Cádiz genetic groups exhibited shorter flower traits (e.g. calix length always <4 mm, except for population D04 from Algarve; **Table S3**). By contrast, individuals of the Tetraploid group tended to exhibit significantly larger floral traits (**Figure 3; Table S2**). However, it is important to note that flowers with small floral traits (e.g. calix < 4mm) were also observed in most populations of the Tetraploid group. In some tetraploid populations (e.g. M13), mean floral traits were even comparable to those of diploid populations from the Cádiz and Algarve groups (**Table S2**).

**Table 1.**
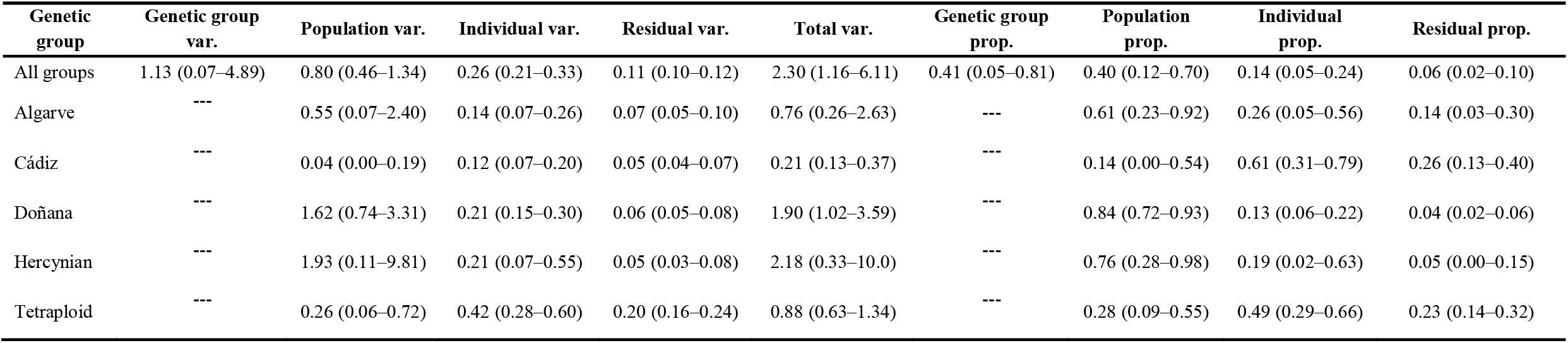
Bayesian hierarchical variance partitioning of calyx size across genetic groups. For the full dataset (“All groups”), variance (var.) components for the genetic group, population, individual, and residual levels are shown, as estimated from a hierarchical model fitted with brms. For each genetic group analyzed separately (Algarve, Cádiz, Doñana, Hercynian, and Tetraploid), variance components correspond to population, individual, and residual levels. Values represent posterior means, with 95% credible intervals in parentheses. Total variance is the sum of all estimated components within each model. Proportional (prop.) contributions indicate the fraction of total variance attributable to each hierarchical level.

**Figure 3.**
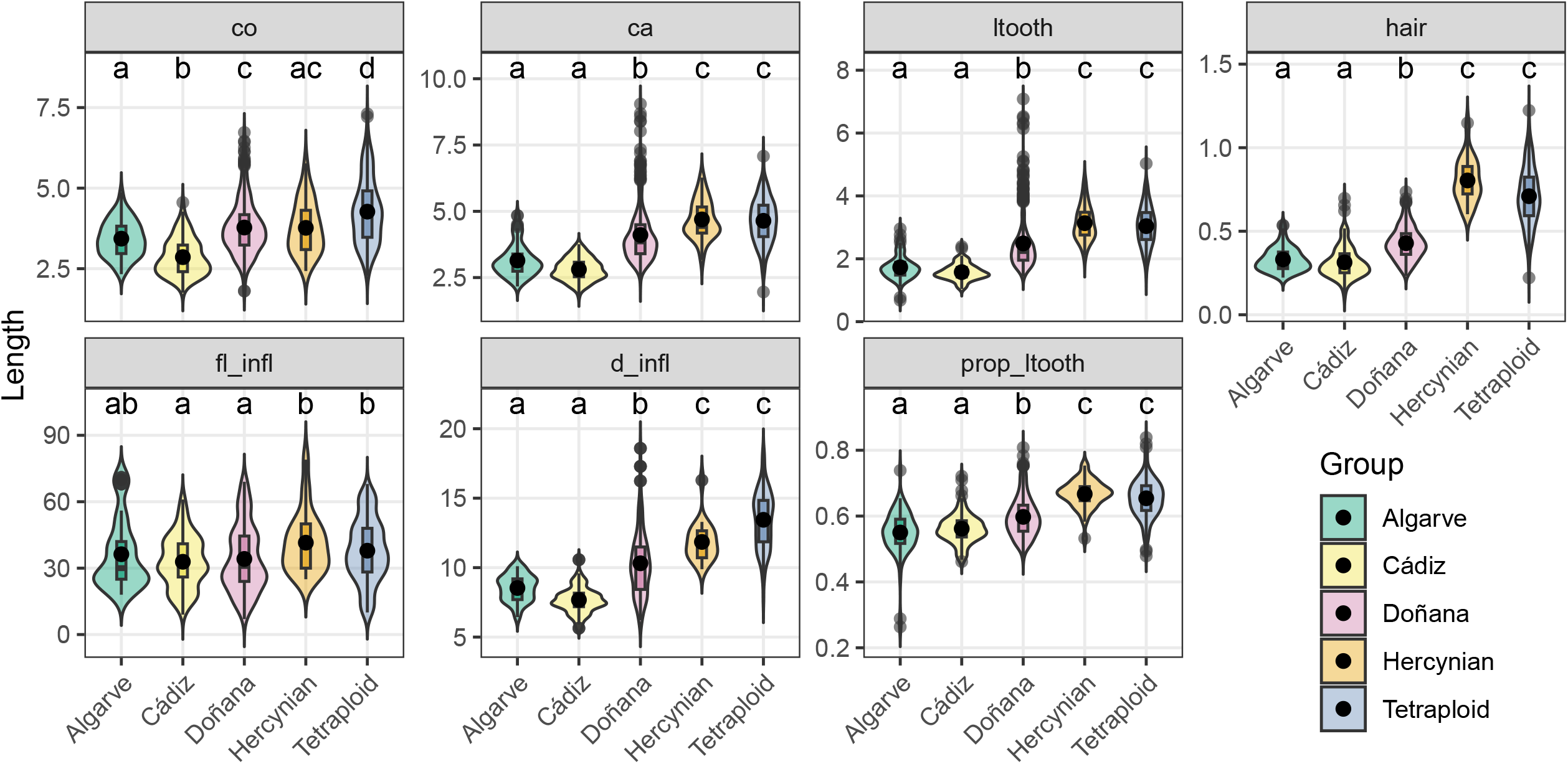
Comparison of morphological traits across genetic groups. Violin plots combined with boxplots show the full distribution of values obtained for each measured morphological trait: corolla length (co), calyx length (Ca), length of the longest calyx tooth (ltooth), length of the longest calyx hair (hair), number of flowers in the inflorescence (fl_infl), inflorescence diameter (d_infl), and proportion of the calyx tooth (prop_ltooth). All metrics are in millimeters. Boxplots represent the interquartile range (IQR); the thin black line indicates 1.5× the IQR, the central line shows the median, and the black dot represent the mean sample value; black dots outside the IQR represent the outlayer values. Different letters above violin plots represent significant pairwise differences among genetic groups according to post-hoc tests on the Dunn test (adjusted p < 0.05).

Interestingly, the diploid Hercynian group also tended to exhibit significantly larger floral traits, comparable to those of the Tetraploid group (**Figure 3; Table S2**), while individuals from populations of Doñana genetic group display the highest variability in most of the floral traits (i.e. co, ca, ltooth, and d_infl), and a broad overlap with all other groups, with some populations with individuals similar in size to the smaller-flowered Algarve and Cádiz groups, others with intermediate characters (e.g. calix length between 4 and 5 mm) and still others with large flower traits resembling those of the Tetraploid and Hercynian genetic groups. One population of western range within the Doñana group (M03) exhibited the largest calyx sizes among all populations (**Table S3**). In fact, Doñana group was the one that exhibited higher percentage of variance attributable to differences among populations (85%; **Table 1**). In contrast, populations from Cádiz genetic group showed the lower percentage of variance attributable to differences among populations (14%), and most of variance (61%) in this group where within individuals; still, variance within Cádiz individuals was the lowest observed in any genetic group (mean=0.12; **Table 1**).

In line with these morphological trends, the PCA clearly separated the Tetraploid and Hercynian clusters from the diploid Algarve and Cádiz genetic groups. However, individuals from Doñana showed a broad overlap with all other groups, further illustrating the high degree of morphological variability across genetic groups (**Figure S3**).

### Soil variables

While tetraploid and diploid populations showed significant differences in all measured soil variables, they also show overlaps in all values (**Figure 4**). Tetraploid populations showed higher values of pH, EC, OM, WRP, and WRC, than diploid populations. When analysing soil variables by genetic groups, only WRC showed significant differences among the diploid genetic groups (i.e. Algarve, Cádiz, Doñana, and Hercynian). Among the four groups, the Hercynian populations generally exhibited higher values, statistically similar to tetraploid populations for most soil variables (i.e. EC, OM, and WRC). The only variable for which the Hercynian group was statistically similar to the other diploid groups but differed significantly from tetraploids was WRP.

**Figure 4.**
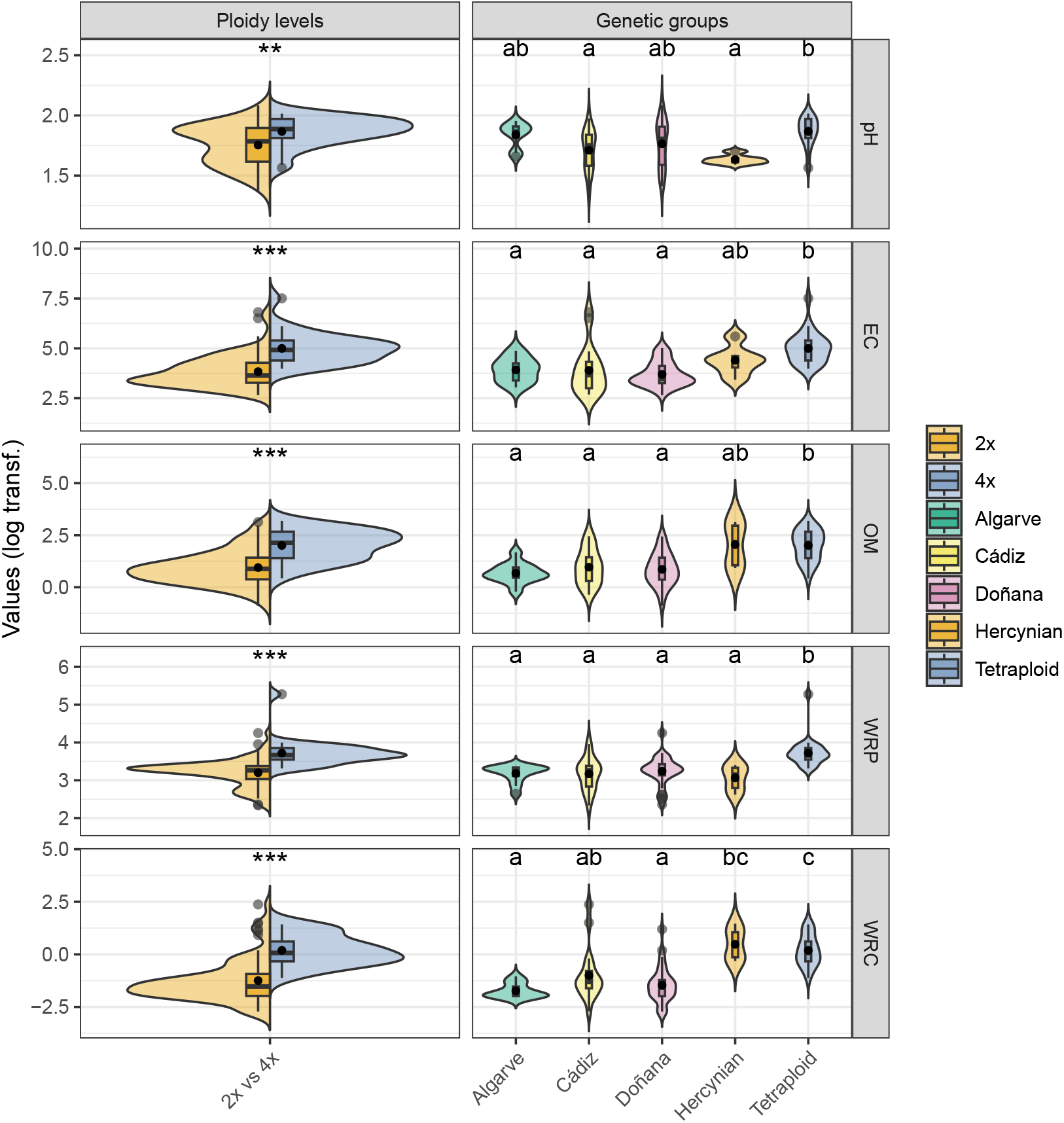
Comparison of soil characteristics across ploidy levels and genetic groups. Violin plots show the distribution of values for diploid (2x) and tetraploid (4x) cytotypes (left column) and for the main genetic groups (right column: Algarve, Cádiz, Doñana, Hercynian and Tetraploid). Each panel corresponds to a different trait electrical conductivity (EC), content in organic matter (OM), pH, water retention potential (WRP), and water retention capacity (WRC). Boxplots represent the interquartile range (IQR); the thin black line indicates 1.5× the IQR, the horizontal central line shows the median, and the black dot within the boxplot represent the mean; black dots outside the IQR represent the outlayer values. Asterisks denote significant differences between ploidy levels (** p < 0.01). Different letters above violin plots represent significant pairwise differences among genetic groups according to post-hoc tests on the Dunn test (adjusted p < 0.05) p < 0.05).

For pH, the Algarve and Doñana groups had the highest variability, together with the Tetraploid populations, whereas Hercynian showed the lowest values. A considerable overlap among both ploidy levels and genetic groups was evident in the PCA analysis (**Figure S4**). Soil EC, OM, and WRP align with axis 1 of the PCA, pH aligns with axis 2, and WRC aligns between the two axes. The first two components of the PCA represented the 56.4% and 21.2 % of the total variation, respectively. The populations displayed in the biplot of these two axes show consistent overlap among ploidy levels and genetic groups.

### Bioclimatic variables

When analysing bioclimatic variables through the first five axes of a PCA (**Figure 5**), those axes represented 37.1, 26.9, 12.2, 10.0, and 4.1% of the variation, respectively. We found significant differences between ploidy levels for axis 1 of the PCA (PC1), although some overlap was observed, mainly due to the Hercynian diploid populations and the high variability within the Tetraploid group (**Figure S6**). Furthermore, when analysing the bioclimatic variables across the genetic groups, we found no PCA axis showing significant differences, except for Tetraploid on axis 1 (PC1), and between Algarve and Doñana on axis 2 and 3 (PC2 and PC3) (**Figure 5**). The biplot of the populations in the two first axes of the PCA show these overlaps more clearly. Populations spread through the first axis of the PCA, with the genetic groups Algarve, Cádiz, and Doñana clustering in the left of the biplot, whereas the two populations of the Hercynian group spreading towards the right. The Tetraploid group overlap extensively with Doñana and Hercynian populations (**Figure S6**).

**Figure 5.**
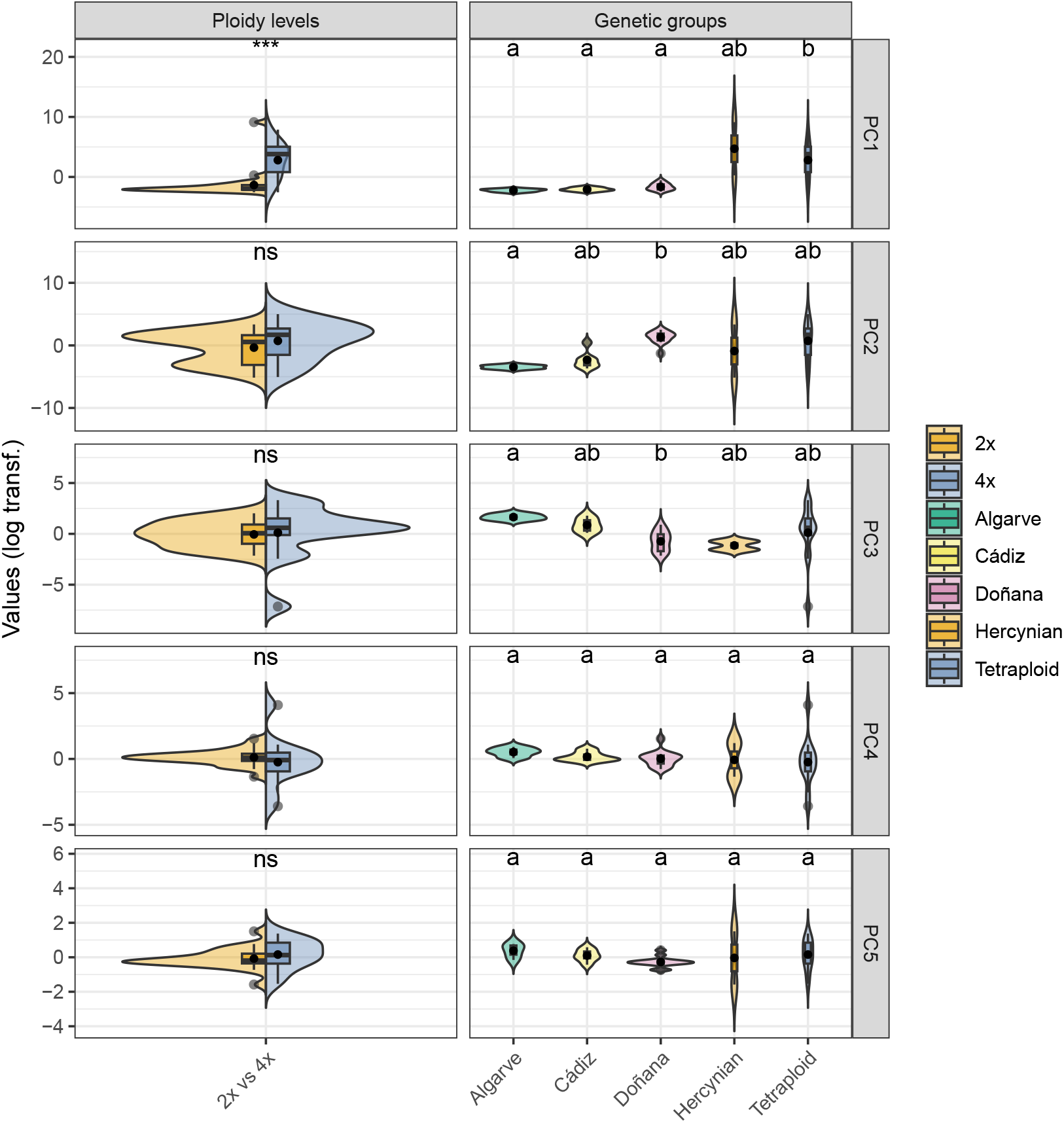
Comparison of bioclimatic variables across ploidy levels and genetic groups. Violin plots show the distribution of values for diploid (2x) and tetraploid (4x) cytotypes (left column) and for the main genetic groups (right column: Algarve, Cádiz, Doñana, Hercynian and Tetraploid). Each panel corresponds to a different principal component (i.e., from PC1 to PC5). Boxplots represent the interquartile range (IQR); the thin black line indicates 1.5× the IQR, the horizontal central line shows the median, and the black dot within the boxplot represent the mean; black dots outside the IQR represent the outlayer values. Asterisks denote significant differences between ploidy levels (***, p < 0.001; ns, non-significant). Different letters above violin plots represent significant pairwise differences among genetic groups according to post-hoc tests on the Dunn test (adjusted p < 0.05).

### Ecological communities

The composition of the plant communities associated with the studied populations showed no clear differentiation across ploidy levels, as revealed by the PCoA based on Hellinger-transformed species cover. The first two PCoA axes explained 21.0% and 12.7% of the variation in community composition, respectively. Jointly both axes do not separate diploid and tetraploid populations (**Figure 6**), with diploid populations overlapping entirely the tetraploid populations. When examined at the level of genetic groups, Algarve, Cádiz, and Doñana populations formed overlapping but distinguishable clusters, whereas the Hercynian and Tetraploid genetic groups were clearly isolated from the remaining groups, indicating greater compositional distinctiveness (**Figure 6**).

**Figure 6.**
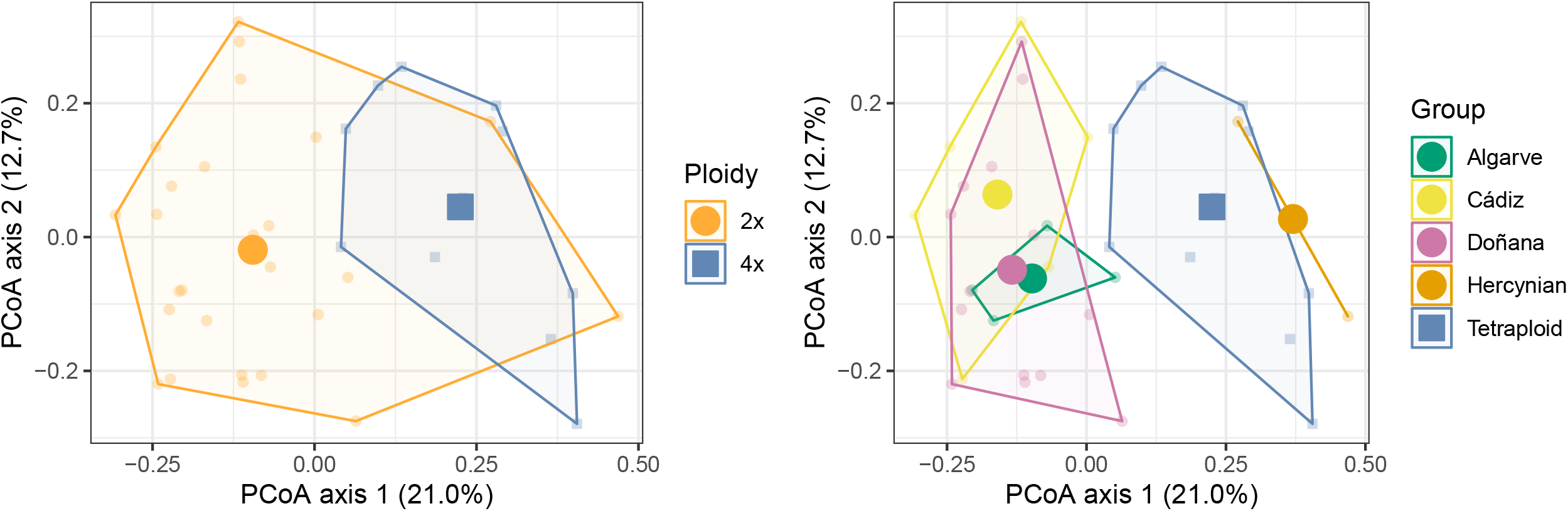
Principal Coordinates Analysis (PCoA) of vegetation dissimilarity across ploidy levels and genetic groups for populations of Thymus sect. Mastichina. PCoA ordination of samples coloured by ploidy level (diploid 2x, tetraploid 4x; left plot) and genetic groups (Algarve, Cádiz, Doñana, Hercynian and Tetraploid; right plot). Convex hulls and centroids summarize the multivariate dispersion and central tendency for each group. Circles and squares represent diploid and tetraploid populations, respectively.

Hercynian populations exhibited the lowest species richness in their communities (8 species), whereas the rest of the groups showed much higher richness, varying from 29 to 39 (mean 35.25). *Pistacia lentiscus* was the only species that appeared in communities of the four genetic groups. No single species was exclusive of the communities of any particular *Thymus* genetic group. Multiple species appeared in communities of both tetraploid and most diploid genetic groups (e.g. *Lavandula pedunculata*, missing only from Cadiz group; *Pinus pinea, Chamaerops humilis, Rhamnus alaternus, or Cistus salviifolius*, missing only from Hercynian group). However, communities of the southwestern diploid genetic groups (Algarve, Doñana and Cádiz) shared some characteristic species (e.g., *Cistus libanotis, Halimium commutatum* or *Halimium halimifolium*).

### Linking genetic ancestry to morphological traits

Our comparison between mixed models with and without sex as a random effect resulted in lower AICc values for the models without sex as random effect for all cases but for those on the corolla length (**Table 2 and Figure S6**). Hence, all reported models are linear models without sex but those for the corolla length (**Table 2**).

**Table 2.** Coefficients and significance levels of univariate and multivariate models between morphological traits and percentage of genetic ancestry (groups 1-5) for individuals in each of the five genetic groups (Algarve, Cádiz, Doñana, Hercynian, and Tetraploid; columns). All models correspond with linear models but those on the corolla that were fit using linear mixed models accounting for the sex of the individual as a random effect. Coefficients and p-values that are under the significance level (p < 0.05) are indicated in bold. In multivariate models, the ancestry corresponding to the group that is analysed are kept out of the models (cells in blank).

|  |  | Univariate models |  |  |  |  |  |  |  |  |  | Multivariate models |  |  |  |  |  |  |  |  |  |
| --- | --- | --- | --- | --- | --- | --- | --- | --- | --- | --- | --- | --- | --- | --- | --- | --- | --- | --- | --- | --- | --- |
|  |  | Algarve<br>(n=23) |  | Cádiz<br>(n=21) |  | Doñana<br>(n=67) |  | Hercynian<br>(n=8) |  | Tetraploid<br>(n=55) |  | Algarve<br>(n=23) |  | Cádiz<br>(n=21) |  | Doñana<br>(n=67) |  | Hercynian<br>(n=8) |  | Tetraploid<br>(n=55) |  |
| Predictors |  | Est. | p | Est. | p | Est. | p | Est. | p | Est. | p | Est. | p | Est. | p | Est. | p | Est. | p | Est. | p |
| Calix | (Intercept) |  |  |  |  |  |  |  |  |  |  | **3.09** | **<0.001** | **2.83** | **<0.001** | **3.58** | **<0.001** | **5.55** | **<0.001** | **4.96** | **<0.001** |
|  | group1 (Tetraploid) | -0.01 | 0.398 | -0.02 | 0.063 | 0.02 | 0.369 | **-0.03** | **0.026** | 0.02 | 0.117 | -0.00 | 0.685 | -0.02 | 0.159 | -0.00 | 0.976 | -0.02 | 0.387 |  |  |
|  | group2 (Hercynian) | 0.06 | 0.085 | 0.01 | 0.821 | **0.19** | **<0.001** | **0.02** | **0.007** | -0.04 | 0.069 | 0.06 | 0.188 | -0.01 | 0.819 | **0.13** | **<0.001** |  |  | -0.04 | 0.087 |
|  | group3 (Algarve) | 0.01 | 0.611 | 0.13 | 0.108 | **0.11** | **<0.001** | -0.04 | 0.082 | 0.01 | 0.761 |  |  | 0.09 | 0.227 | **0.06** | **0.001** | -0.01 | 0.495 | 0.01 | 0.766 |
|  | group4 (Doñana) | -0.02 | 0.823 | 0.02 | 0.082 | **-0.04** | **<0.001** | 0.02 | 0.626 | 0.00 | 0.977 | -0.00 | 0.992 | 0.01 | 0.274 |  |  | 0.00 | 0.996 | -0.01 | 0.880 |
|  | group5 (Cádiz) | -0.06 | 0.632 | -0.00 | 0.748 | **-0.08** | **0.028** | **-0.09** | **0.012** | -0.04 | 0.342 | -0.02 | 0.870 |  |  | -0.03 | 0.192 | -0.06 | 0.230 | -0.03 | 0.452 |
| Corolla | (Intercept) |  |  |  |  |  |  |  |  |  |  | **3.46** | **<0.001** | **2.82** | **<0.001** | **3.36** | **<0.001** | **3.70** | **<0.001** | **4.44** | **<0.001** |
|  | group1 (Tetraploid) | -0.01 | 0.240 | -0.01 | 0.746 | 0.00 | 0.922 | 0.03 | 0.090 | 0.01 | 0.654 | -0.01 | 0.090 | 0.00 | 1.000 | -0.00 | 0.561 | **0.04** | **0.002** |  |  |
|  | group2 (Hercynian) | 0.02 | 0.607 | -0.04 | 0.449 | **0.10** | **<0.001** | -0.01 | 0.192 | -0.03 | 0.328 | 0.01 | 0.765 | -0.06 | 0.141 | **0.08** | **<0.001** |  |  | -0.04 | 0.151 |
|  | group3 (Algarve) | 0.01 | 0.280 | 0.16 | 0.088 | **0.06** | **<0.001** | 0.03 | 0.238 | 0.03 | 0.528 |  |  | 0.09 | 0.194 | **0.03** | **0.015** | 0.02 | 0.187 | 0.05 | 0.288 |
|  | group4 (Doñana) | -0.04 | 0.558 | **0.03** | **0.026** | **-0.02** | **0.002** | -0.04 | 0.414 | 0.01 | 0.794 | -0.01 | 0.856 | **0.03** | **0.003** |  |  | 0.01 | 0.793 | 0.01 | 0.765 |
|  | group5 (Cádiz) | 0.14 | 0.201 | -0.02 | 0.109 | -0.04 | 0.166 | 0.01 | 0.801 | -0.01 | 0.840 | 0.05 | 0.524 |  |  | 0.01 | 0.694 | **-0.07** | **0.009** | -0.00 | 0.956 |
| Diameter infl. | (Intercept) |  |  |  |  |  |  |  |  |  |  | **8.27** | **<0.001** | **7.48** | **<0.001** | **9.58** | **<0.001** | **12.91** | **0.002** | **13.97** | **<0.001** |
|  | group1 (Tetraploid) | 0.01 | 0.467 | 0.01 | 0.859 | 0.01 | 0.772 | -0.03 | 0.581 | 0.05 | 0.298 | 0.02 | 0.273 | 0.01 | 0.746 | -0.01 | 0.676 | -0.03 | 0.777 |  |  |
|  | group2 (Hercynian) | 0.11 | 0.067 | 0.07 | 0.569 | **0.30** | **<0.001** | 0.01 | 0.645 | -0.07 | 0.237 | 0.11 | 0.083 | 0.06 | 0.669 | **0.22** | **0.003** |  |  | -0.07 | 0.297 |
|  | group3 (Algarve) | -0.01 | 0.388 | -0.08 | 0.676 | **0.16** | **<0.001** | -0.03 | 0.687 | -0.01 | 0.913 |  |  | -0.10 | 0.649 | 0.09 | 0.073 | -0.01 | 0.901 | -0.03 | 0.842 |
|  | group4 (Doñana) | **-0.26** | **0.021** | 0.01 | 0.603 | **-0.06** | **0.002** | 0.04 | 0.738 | 0.06 | 0.537 | -0.21 | 0.051 | 0.02 | 0.592 |  |  | 0.02 | 0.901 | 0.05 | 0.646 |
|  | group5 (Cádiz) | -0.19 | 0.330 | -0.01 | 0.530 | -0.11 | 0.157 | -0.03 | 0.826 | -0.13 | 0.272 | -0.05 | 0.780 |  |  | -0.03 | 0.653 | 0.03 | 0.890 | -0.12 | 0.346 |
| Hair | (Intercept) |  |  |  |  |  |  |  |  |  |  | **0.31** | **<0.001** | **0.28** | **<0.001** | **0.41** | **<0.001** | **0.88** | **0.001** | **0.74** | **<0.001** |
|  | group1 (Tetraploid) | -0.00 | 0.340 | -0.00 | 0.315 | 0.00 | 0.378 | -0.00 | 0.774 | 0.00 | 0.421 | -0.00 | 0.885 | -0.00 | 0.964 | 0.00 | 0.471 | -0.01 | 0.259 |  |  |
|  | group2 (Hercynian) | **0.01** | **0.001** | 0.00 | 0.827 | **0.01** | **0.047** | -0.01 | 0.319 | -0.00 | 0.398 | **0.01** | **0.006** | -0.00 | 0.454 | 0.01 | 0.125 |  |  | -0.00 | 0.408 |
|  | group3 (Algarve) | 0.00 | 0.729 | **0.04** | **0.004** | 0.00 | 0.156 | 0.00 | 0.747 | 0.00 | 0.697 |  |  | **0.03** | **0.002** | 0.00 | 0.916 | 0.00 | 0.586 | 0.00 | 0.684 |
|  | group4 (Doñana) | -0.00 | 0.730 | **0.01** | **<0.001** | -0.00 | 0.079 | 0.00 | 0.217 | -0.00 | 0.614 | 0.00 | 0.859 | **0.01** | **<0.001** |  |  | -0.00 | 0.978 | -0.00 | 0.586 |
|  | group5 (Cádiz) | -0.01 | 0.289 | **-0.00** | **0.011** | -0.00 | 0.878 | -0.01 | 0.073 | -0.00 | 0.772 | -0.00 | 0.689 |  |  | 0.00 | 0.779 | -0.00 | 0.907 | -0.00 | 0.827 |
| L. tooth | (Intercept) |  |  |  |  |  |  |  |  |  |  | **1.70** | **<0.001** | **1.58** | **<0.001** | **1.97** | **<0.001** | **3.69** | **<0.001** | **3.24** | **<0.001** |
|  | group1 (Tetraploid) | -0.00 | 0.485 | -0.01 | 0.182 | 0.01 | 0.428 | -0.02 | 0.065 | 0.01 | 0.224 | -0.00 | 0.755 | -0.01 | 0.350 | -0.00 | 0.758 | -0.01 | 0.679 |  |  |
|  | group2 (Hercynian) | 0.04 | 0.103 | 0.00 | 0.982 | **0.16** | **<0.001** | **0.02** | **0.026** | -0.02 | 0.152 | 0.04 | 0.218 | -0.01 | 0.755 | **0.11** | **<0.001** |  |  | -0.02 | 0.195 |
|  | group3 (Algarve) | 0.00 | 0.704 | **0.10** | **0.041** | **0.09** | **<0.001** | -0.03 | 0.089 | 0.01 | 0.805 |  |  | 0.08 | 0.100 | **0.05** | **<0.001** | -0.01 | 0.519 | 0.01 | 0.853 |
|  | group4 (Doñana) | -0.01 | 0.908 | 0.01 | 0.135 | **-0.04** | **<0.001** | 0.03 | 0.450 | 0.01 | 0.677 | 0.01 | 0.914 | 0.01 | 0.363 |  |  | 0.01 | 0.676 | 0.01 | 0.817 |
|  | group5 (Cádiz) | -0.06 | 0.503 | -0.00 | 0.624 | **-0.06** | **0.048** | **-0.07** | **0.020** | -0.04 | 0.233 | -0.03 | 0.721 |  |  | -0.02 | 0.340 | -0.05 | 0.272 | -0.03 | 0.322 |
| Prop. L. tooth | (Intercept) |  |  |  |  |  |  |  |  |  |  | **0.55** | **<0.001** | **0.56** | **<0.001** | **0.56** | **<0.001** | **0.66** | **<0.001** | **0.65** | **<0.001** |
|  | group1 (Tetraploid) | 0.00 | 0.961 | 0.00 | 0.605 | 0.00 | 0.796 | -0.00 | 0.855 | -0.00 | 0.567 | 0.00 | 0.854 | 0.00 | 0.599 | -0.00 | 0.309 | 0.00 | 0.483 |  |  |
|  | group2 (Hercynian) | 0.00 | 0.479 | -0.00 | 0.753 | **0.01** | **<0.001** | 0.00 | 0.686 | 0.00 | 0.634 | 0.00 | 0.553 | -0.00 | 0.883 | **0.01** | **<0.001** |  |  | 0.00 | 0.516 |
|  | group3 (Algarve) | -0.00 | 0.832 | 0.01 | 0.283 | **0.00** | **<0.001** | -0.00 | 0.385 | 0.00 | 0.922 |  |  | 0.01 | 0.288 | **0.00** | **0.001** | -0.00 | 0.637 | -0.00 | 0.934 |
|  | group4 (Doñana) | 0.00 | 0.881 | -0.00 | 0.902 | **-0.00** | **<0.001** | 0.00 | 0.230 | 0.00 | 0.160 | 0.00 | 0.768 | -0.00 | 0.870 |  |  | 0.00 | 0.358 | 0.00 | 0.197 |
|  | group5 (Cádiz) | -0.01 | 0.521 | -0.00 | 0.733 | -0.00 | 0.387 | -0.00 | 0.383 | -0.00 | 0.234 | -0.01 | 0.676 |  |  | 0.00 | 0.622 | -0.00 | 0.489 | -0.00 | 0.276 |

Across genetic groups, the models revealed marked differences in the extent to which percentage of genetic ancestry affect floral morphological traits. Overall, Doñana individuals suffered the strongest and most consistent effect of percentage of genetic ancestry on floral morphology: individuals from this group exhibited significant relationships between ancestry from other diploid groups and most major traits examined (**Table 2**). In particular, the models suggest a positive effect of ancestry proportion from Hercynian and Algarve groups. These effects remain mostly significant when all ancestry groups are combined in multivariate models, indicating a strong and independent effect.

In contrast, Tetraploid individuals exhibited nonsignificant associations in all cases (any combination of morphological trait and ancestry group). This lack of effects is evident in both univariate and multivariate models (**Table 2**). The rest of the genetic groups displayed only sporadic and trait-specific associations, and most of them changed from significant effects in univariate models to nonsignificant in multivariate models (e.g. the negative effect of Tetraploid ancestry in calyx length for the Hercynian individuals). The only effects that are significant both in univariate and multivariate models are: a) a negative effect of percentage of Hercynian ancestry on the length of the longest hair of the calyx tooth (hair) for Algarve individuals, b) the positive effect of percentage of Doñana ancestry on corolla length (co) and length of the hair for Cádiz individuals, and c) the positive effect of percentage of Algarve ancestry on the length of the hair for Cádiz individuals. There are only two effects that appear in multivariate models that were nonsignificant in univariate models: a negative effect of percentage of Cádiz ancestry and a positive effect of percentage of Tetraploid ancestry on the corolla length for the Hercynian populations. This was the only effect detected for Tetraploid ancestry in any diploid population, and although it was statistically significant, the estimated effect size was very small, indicating a weak signal.

## Discussion

Our results reveal a remarkable level of intraspecific complexity within *Thymus* sect. *Mastichina*, characterized by substantial overlap among genetic lineages at the morphological and ecological levels. While some degree of differentiation was observed between diploid and tetraploid individuals, the high variability complicates taxonomic delimitation based on discrete morphological traits. This discordance between phenotypic variation and genetic structure likely reflects a combination of phenotypic plasticity, recent divergence, or incomplete reproductive isolation (Villellas et al. 2014; Castillo et al. 2018). Taken together, these findings suggest that *Thymus* sect. *Mastichina* constitutes a complex of cryptic lineages, where genetic differentiation is not consistently mirrored by phenotypic or ecological distinctiveness.

### Ginodioecy and sexual dimorphism

In our study, the only trait showing significant differences associated with sex was corolla size, with hermaphrodite flowers exhibiting larger corollas than females. This pattern of floral dimorphism is among the most frequently reported floral traits exhibiting sexual dimorphism in angiosperms, including other *Thymus* species (Thompson et al. 2002; StakelienE and LožienE 2014). This is often attributed to developmental associations between the corolla and stamens, as well as differing selective pressures: larger corollas in pollen-producing flowers enhance pollinator attraction and male fitness, while smaller corollas in female flowers may reflect resource allocation to seed production or reduced need for pollinator attraction (Delph et al. 1996; Niu et al. 2015).

### Morphological variability

The morphological characterization of *Thymus* sect. *Mastichina* reveal significant differences primarily associated with ploidy level. As expected, tetraploid individuals exhibit, on average, larger floral traits than their diploid counterparts, a pattern commonly attributed to polyploidy-induced increases in cell size and, consequently, in the size of reproductive organs (Levin 1983; Ramsey and Ramsey 2014). However, this trend is not consistent across all diploid populations, since individuals from Hercynian and from some populations from the Doñana genetic group displayed consistently larger floral traits, comparable to those of the Tetraploid group. This contrasts with the smaller morphological features of diploid individuals from other southwestern populations, particularly those within the Cádiz, and Algarve genetic groups. Otherwise, in all populations of the Tetraploid group we found some flowers with size comparable to the smaller flowers in the Cádiz and Algarve groups. This marked morphological overlap between certain tetraploid and diploid individuals suggests that, although ploidy level can influence floral size (Landis et al. 2020; Vilcherrez-Atoche et al. 2022; García-Muñoz et al. 2023), complex interplay of genetic (Weiss et al. 2005; Krizek and Anderson 2013), ecological (Chen et al. 2025; Day Briggs and Anderson 2025), or ontogenic (Diggle 1997; Bull-Hereñu and Ronse De Craene 2020) factors may also shape trait expression.

Although we hypothesized that interploidy genetic introgression could be influencing observed morphological traits, we found no evidence that introgression with the Tetraploid group is affecting diploid’s morphology. Tetraploid introgression within the diploid groups did not have a significant impact on the studied morphological variables but for the corolla length in Hercynian individuals. In contrast, introgression among diploid genetic groups seems to significantly affect floral morphology. Namely, ancestry proportion mainly from Hercynian and, to a lesser extent, from Algarve lineages seem to affect floral size in Doñana individuals. Thus, introgression from these groups could be modulating morphological traits in Doñana individuals, potentially contributing to the high observed phenotypic variability in this genetic group. García-Cárdenas *et al.*(2025) postulated that Doñana group is likely the most recently established, more probably from the adjacent groups of Cádiz or Algarve. Nevertheless, admixture with the Hercynian group appears to be responsible for the longer morphological traits, particularly calyx length and tooth length, observed in this group, which may explain, at least partially, the resulting pattern of morphological overlap.

### Ecological niches

The analysis of environmental conditions revealed significant differences between ploidy levels, with the tetraploid cluster occupying a distinct ecological niche relative to the diploid groups (with some exceptions, see below). These findings suggest niche differentiation associated primarily with ploidy level rather than among diploid lineages, consistent with previous studies highlighting the ecological significance of polyploidy (e.g., Ramsey 2011; Soltis et al. 2014). However, the diploid Hercynian group partially overlaps with the Tetraploid group in both geographic distribution and environmental conditions. In this context, García-Cárdenas *et al*. (2025) suggested that the Hercynian diploids may represent a relict lineage derived from an ancestral diploid population that has been largely replaced by tetraploids throughout much of its range. Meanwhile, the southwestern Iberian Peninsula may have functioned as a Pleistocene glacial refugium, offering long-term geographic and ecological stability that facilitated the persistence and divergence of isolated lineages, promoting speciation and the retention of distinctive genetic and morphological traits in the diploid taxa present in this region (Girón et al. 2012).

### Taxonomic and conservation implications

The limited morphological resolution observed across populations of this section underscores a broader challenge in plant systematics: the frequent mismatch between phenotype and underlying genetic structure, especially in recently diverged or rapidly evolving groups (Bickford et al. 2007; Struck et al. 2018). As humans, we depend strongly on visual information to interpret the natural world, and classical approaches to species classification have therefore been based mainly on morphological traits (Bickford et al. 2007; Hending 2025). However, in taxonomically complex groups, the limited diagnostic power of morphological traits highlights the challenges of using phenotypic traits alone for species delimitation. Addressing this issue ideally requires the integration of multiple lines of evidence to clarify species boundaries and support appropriate conservation actions (Federici et al. 2013; Fišer et al. 2018). Molecular approaches, such as DNA Barcoding has proven to be an effective tool for species identification (Antil et al. 2023) and are increasingly applied to detect cryptic species (e.g. Nieto-Lugilde et al. 2024). Genome size has also emerged as a reliable character for species delimitation (Kron et al. 2007; Fomicheva and Domblides 2023). Additionally, integrative taxonomy considers other sources of information, such as geographic distribution and niche differentiation, which can further contribute to robust species delimitation (Dayrat 2005; De Luna-Bonilla et al. 2024; Wang et al. 2025). Previous genomic work in *Thymus* sect. *Mastichina* (García-Cárdenas et al. 2025) provides a useful foundation for developing DNA barcoding tools, although additional research will be required to fully achieve this goal. For species differentiation of our study system, ploidy level constitutes a more reliable character for species identification than flower morphology and, therefore, we advocate its incorporation into taxonomic assessments. Flow cytometry is widely used for estimating plant genome sizes and is an accessible method that can be conducted in external specialized laboratories at relatively low cost (Galbraith et al. 2021; Sliwinska et al. 2022). Thus, following Girón *et al*. (2012) and García-Cárdenas *et al*. (2025), we can stablish a reliable distinction between species within *Thymus* sect *Mastichina*: diploid individuals of with 2C DNA content from 1.32 to 1.84 pg should be considered as *T. albicans* Hoffmanns. & Link, whereas tetraploids with 2C DNA values from 2.31 to 3.07 pg should be considered as *T. mastichina* L.

Although we detected a considerable overlap in ecological niches, this pattern was mainly driven by diploid populations inhabiting subcoastal areas (Algarve, Doñana, and Cádiz groups). In contrast, the diploid Hercynian group occupied a distinctive ecological niche more closely resembling that of the Tetraploid group. The Hercynian group is also morphologically and genetically García-Cárdenas et al. (2025) distinct from other diploid lineages. Taken together, these ecological, morphological, and genomic differences indicate that the Hercynian group represents a distinct evolutionary lineage from *T. albicans* and may warrant recognition as a new species *T. hercynica*. This taxon can be considered cryptic with respect to *T. mastichina* given the strong overlap in floral morphology, ecological niche and geographic distribution, nevertheless it is readily distinguishable based on its 2C DNA content. Given that this lineage is currently known from only two populations, we adopt a cautious approach, and formal nomenclatural validation, including typification and diagnosis, will be addressed in future research.

Concerning Doñana genetic group, the extensive variability in floral traits among populations complicates its taxonomic recognition. Many populations within this group were formerly recognized as *T. mastichina* subsp. *donyanae*. However, we found substantial overlap with the other component of the section in terms of morphology. Genomic evidence clearly supports its phylogenetic placement within *T. albicans* (García-Cárdenas et al. 2025). Given the genomic differentiation among these geographically structured lineages (García-Cárdenas et al. 2025), we propose a geographically based subspecific framework that provides a pragmatic solution for documenting intraspecific genetic diversity. Accordingly, diploid populations in SW Iberia between the Guadiana and Guadalquivir rivers would be referred to the combination *Thymus albicans* subsp. *donyanae* (R. Morales) Rivas Mart. (Rivas Martínez 1988). Lastly, populations east of the Guadalquivir River could be referred as *Thymus albicans* subsp. *gaditanum*, and those west of the Guadiana River could be referred as *T. albicans* subsp. *albicans*.

The recognition of the Doñana populations as *Thymus albicans* subsp. *donyanae* would have relevant implications for conservation, as *T. albicans* is an endangered species (Moreno 2008; Buira et al. 2017). Following IUCN criteria (IUCN, 2012) and given the geographically intermediate position of Doñana populations between the Cádiz and Algarve populations (the sole formerly recognized as *T. albicans*) the extent of occurrence (EOO) would remain geographically restricted to SW Iberia and comparable to previous estimates. However, the area of occupancy (AOO) would increase due to the inclusion of additional populations. The main threats affecting *T. albicans*, land-use change particularly associated with coastal urban pressure and greenhouse agriculture, are also present across the range of *T. albicans* subsp. *donyanae*, although a proportion of its populations occur within protected areas. A comprehensive conservation assessment is beyond the scope of the present study and should be addressed in future research, once the taxonomic framework is resolved. Moreover, given the effect of the introgression on flower morphology of some Doñana populations, we would recommend managing them as differentiated conservation units. This would help to safeguard the genetic and morphological integrity of other diploid groups, especially the morphologically homogeneous and genetically distinct Cádiz group. Future assessments must specifically define these relevant genetic units, evaluate current and emerging threats across populations, and consider the evolutionary potential of these distinct groups to inform effective long-term conservation strategies.

## Conclusions

Our integrative analysis indicates that *Thymus* sect. *Mastichina* represents a complex of cryptic evolutionary lineages characterized by extensive morphological overlap, and limited ecological differentiation among diploid groups, despite a marked genetic structuring. Although ploidy level influences floral traits and ecological niches, phenotypic variation is often decoupled from genetic lineages, possibly due to recent divergence and introgression. Contrary to our expectations, introgression among diploid lineages (rather than interploidy introgression) appears to affect floral morphology, particularly contributing to phenotypic overlap in some lineages. These results highlight the limitations of morphology-based taxonomy in this group and emphasize the need for an integrative framework, proposing the use of genome size, genomic information, and geographic distribution for species delimitation.

## Acknowledgements

This study is part of the Conserva3 project (TED2021-130133B-I00) funded by MCIN/AEI/10.13039/501100011033 with funds from the European Union “NextGenerationEU”/PRTR.

Collection permits were obtained from Consejería de Sostenibilidad, Medio Ambiente y Economía Azul, Junta de Andalucía (Spain), Doñana National Park, and from Departamento de Conservação da Natureza e Biodiversidade (Portugal, permits no. 576– 580/REC/2023). In Doñana National Park, logistic and technical support was provided by ICTS-RBD-CSIC, Ministry of Science and Innovation, and co-financed by FEDER Funds.

Authors would like to thank Andrés Melero, as well as the staff from the Doñana National Park, from the Environmental Agency and the Botanical Network of the Andalusian Government (Junta de Andalucía) for their support in population location and sample collection.

Morphology measurements were performed at the facilities of Herbarium Research Services (CITIUS, University of Seville). Part of the costs of those Research Services were funded by 2023 and 2024 calls of VII PPIT of the University of Sevilla. Additional thanks go to Ramón Morales for his advice and expertise regarding the group’s taxonomy.

## Author contributions

RB and DNL conceived the study and established the methodology. DD, JCV, RB, MO, FJL, JLT, DNL, FJGC, EMC and BJW carried out the sampling. DD and DNL obtained and analysed the soil and climatic data. EMC, DDP, DNL and RB obtained and analysed the morphological data. DNL, JCV, and RBP drafted the manuscript. All authors contributed to the final manuscript and approved the submitted version.

