## Supplemental document for "Genomic structure, introgression, niche overlap, and morphological variation challenges species delimitation in Thymus sect. Mastichina"

**Supplementary material**

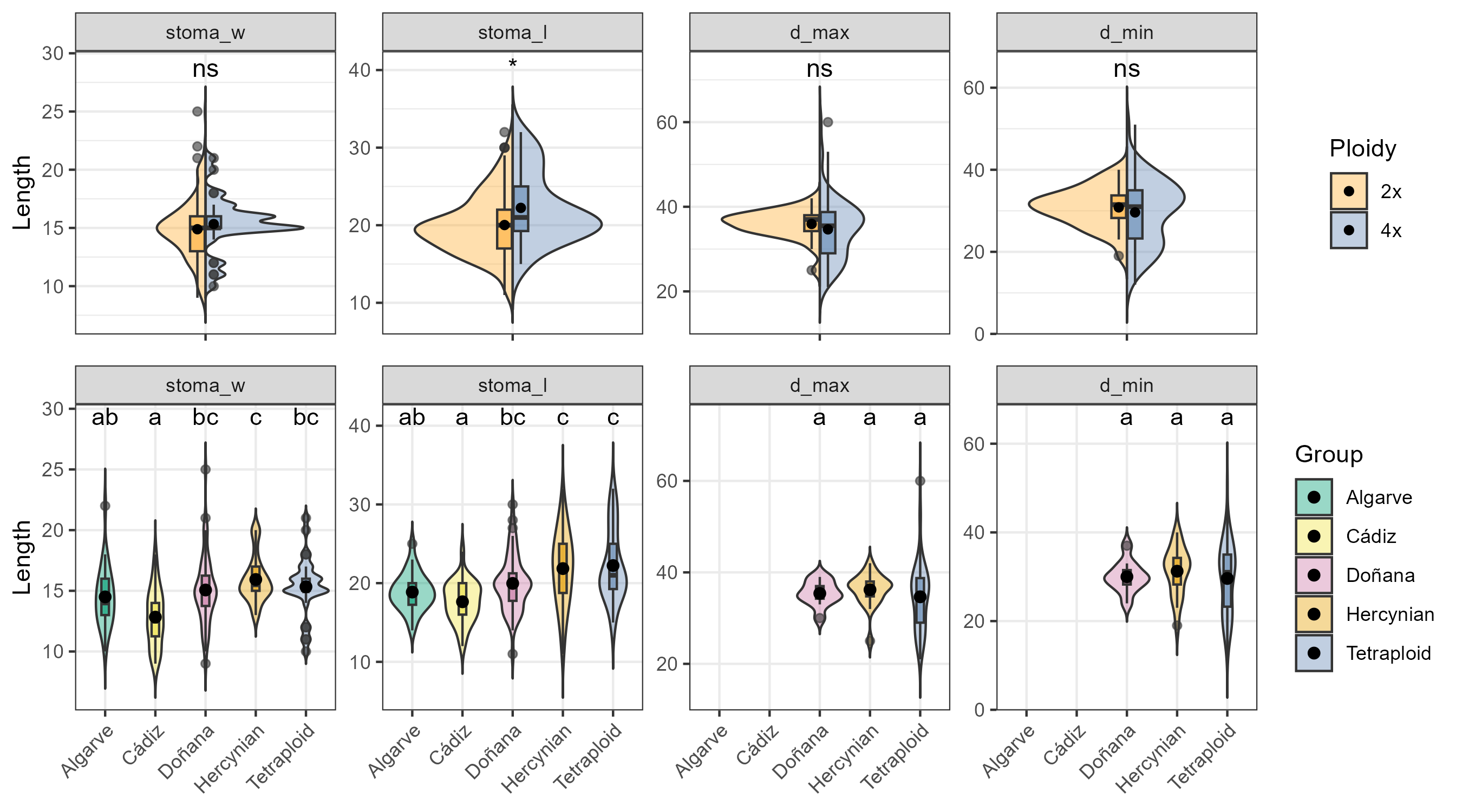

**Figure S1.** *Comparison of morphological traits between ploidy levels and genetic groups for microscopic variables (stoma and pollen grain sizes).* Violin plots combined with boxplots show the full distribution of values obtained for each measured morphological trait: stomata width (stoma_measure_W), stomata length (stoma_measure_L), pollen gran major axis (d_max), and pollen gran minor axis (d_min). All metrics are in micrometres. Boxplots represent the interquartile range (IQR); the thin black line indicates 1.5× the IQR, the central line shows the median, and the black dot represent the mean sample value; black dots outside the IQR represent the outlier values. Asterisks in ploidy level plots denote significant differences between ploidy levels (ns, non-significant; *, P < 0.05; **, *P* < 0.01; and ***, *P* < 0.001). Different letters above figures in genetic group plots represent significant pairwise differences among genetic groups according to post-hoc tests on the Dunn test (adjusted p < 0.05).

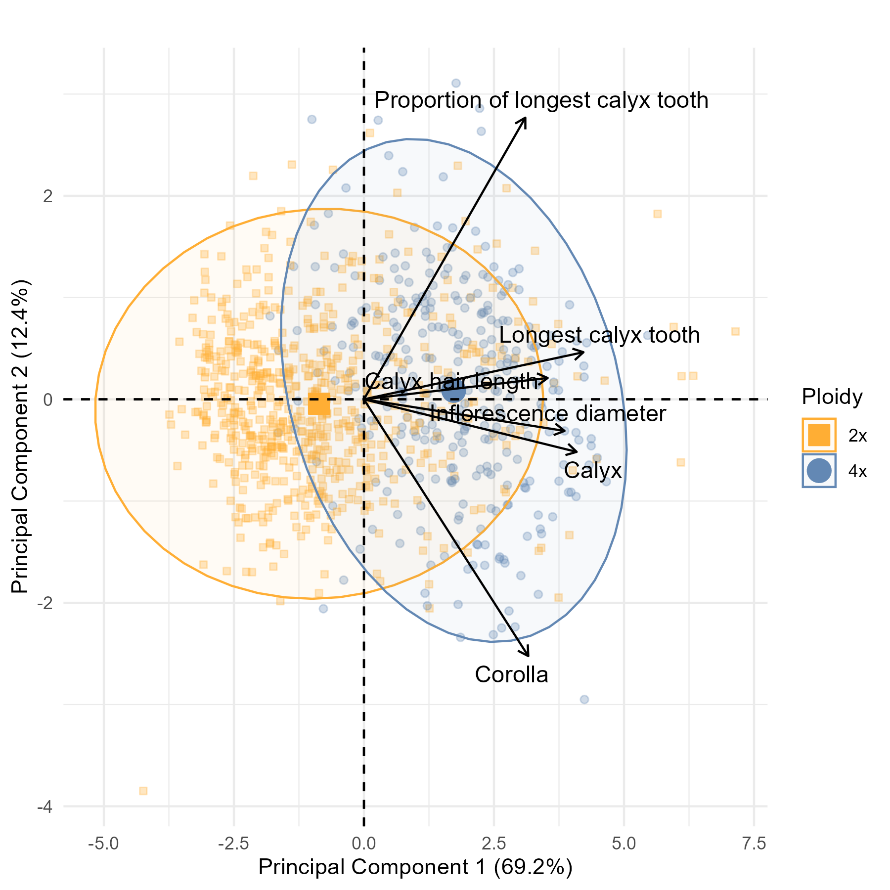

**Figure S2.** *Principal components analysis (PCA) plots depicting variation between diploid (orange squares) and tetraploid (blue circles) individuals based on measurements of the following six floral traits: corolla length, calyx length, length of the longest calyx tooth, length of the longest calyx hair, inflorescence diameter, and proportion of the longest calyx tooth*. Points (squares and circles) represent individual data point, whereas larger symbols indicate the centroids of the 95% confidence ellipses for each genetic cluster.

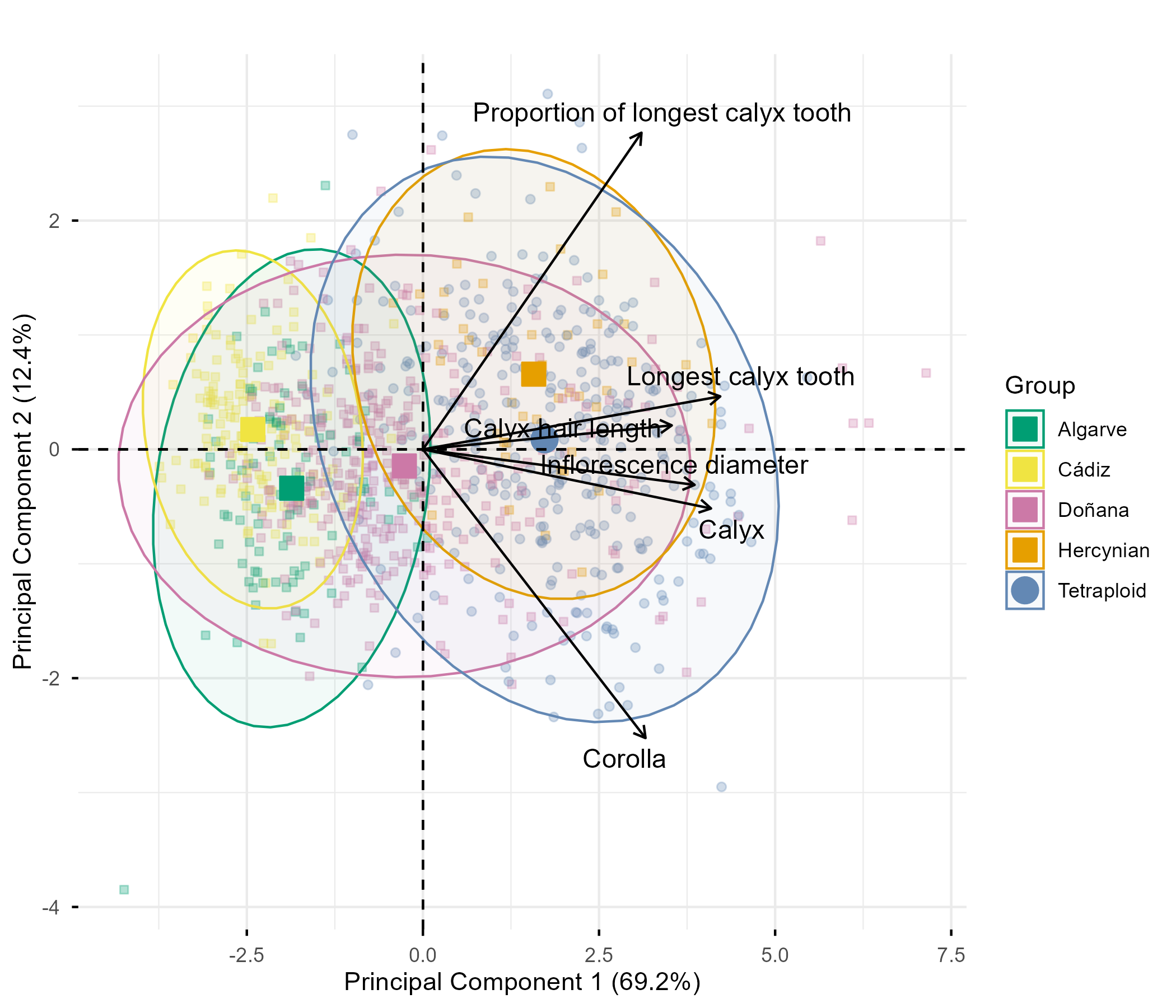

**Figure S3.** *Principal components analysis (PCA) plots depicting variation among the five genetic clusters described in García-Cárdenas et al. (2025): ‘Tetraploid’ (blue), ‘Hercynian’ (orange), ‘Algarve’ (green), ‘Donyana’ (yellow), and ‘Cadiz’ (pink).* Tetraploid individuals are represented by filled circles, while diploid individuals are represented by squares. PCA was based on measurements of the following six floral traits: corolla length, calyx length, calyx hair size, length of the shortest calyx tooth, length of the longest calyx hair, and inflorescence diameter. Smaller symbols represent individual data point, whereas larger symbols indicate the centroids of the 95% confidence ellipses for each genetic cluster.

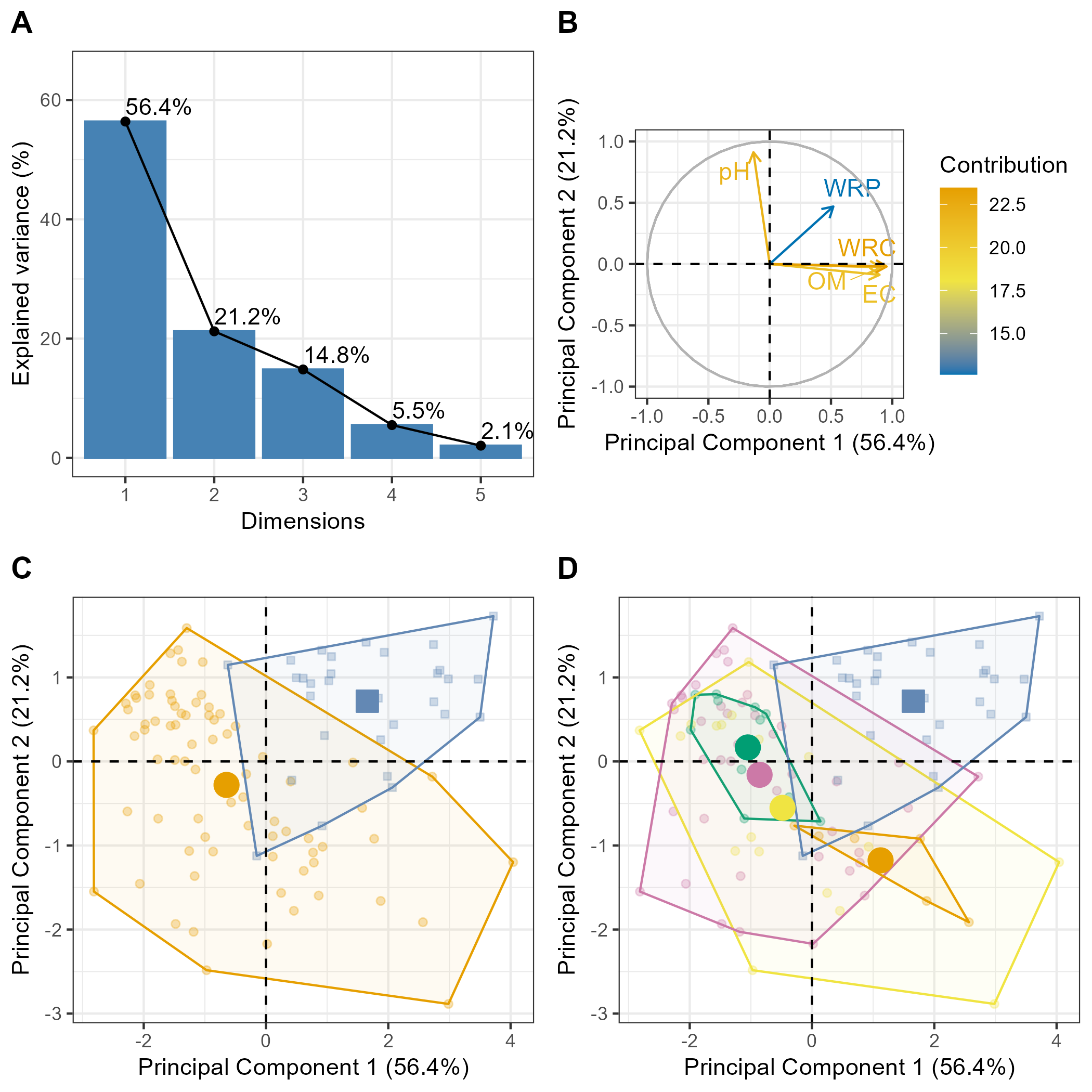

**Figure S4.** Principal Component Analysis (PCA) of soil variables for the samples collected at populations of Thymus Sect. Mastichina across ploidy levels and genetic groups. A, scree plot showing the percentage of variance explained by the first five principal components, with PC1 and PC2 accounting for 56.4% and 21.2% of the total variation, respectively. B, correlation circle displaying the contribution and direction of soil variables to the first two axis of the PCA, with arrow colours indicating variable contributions. C, PCA ordination of samples (populations) in light colours grouped by ploidy level (diploid and tetraploid), showing also group centroids (darker and bigger symbols) and convex hulls enclosing all samples within the same group.D, PCA ordination of samples coloured by genetic group (Algarve, Cadiz, Donyana, Hercynian and Tetraploid), with convex hulls and group centroids.

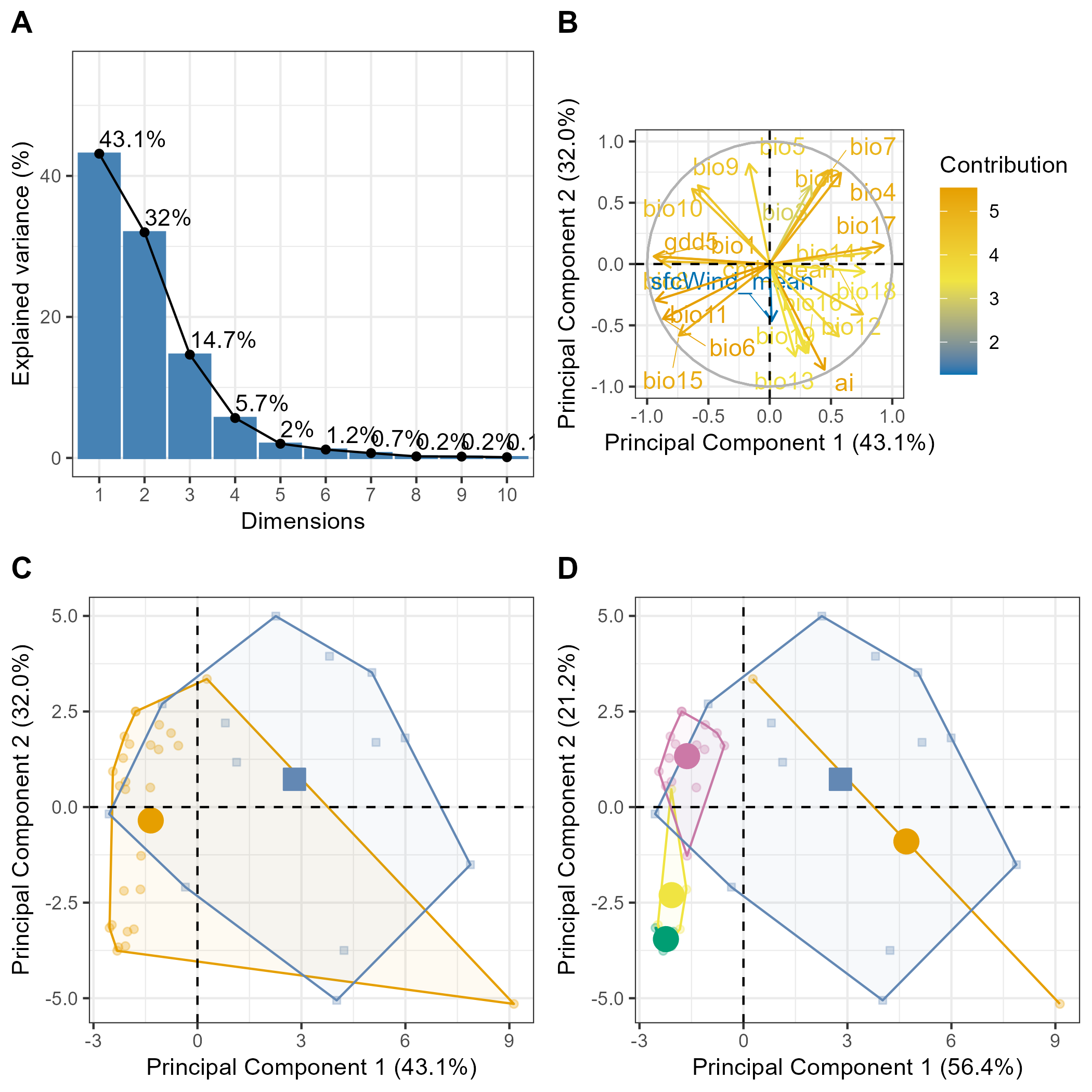
**Figure S5.** Multivariate patterns of climate characteristics based on a Principal Component Analysis (PCA). A, scree plot showing the proportion of variance explained by the first five principal components; PC1 (56.4%) and PC2 (21.2%) together account for most of the variation. B, correlation circle depicting the loadings of bioclimate variables on the first two axis of the PCA, arrow colours indicate variable contributions. C, PCA ordination of samples (i.e., populations) grouped by ploidy level (i.e., between diploid and tetraploid populations), with convex hulls and centroids. D, PCA ordination coloured by genetic group, showing also convex hulls and centroids for each group.

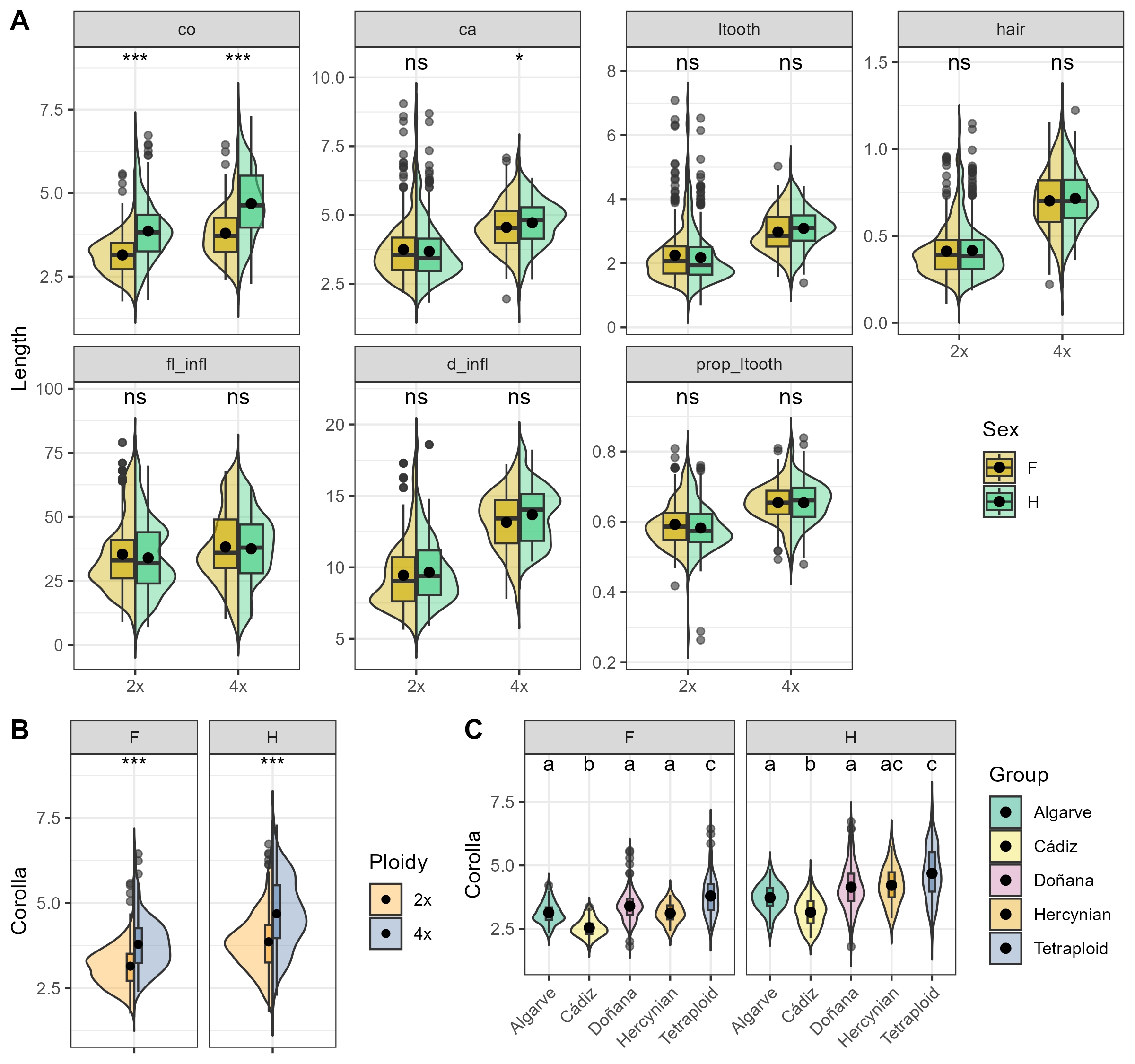

**Figure S6.** *Effect of sex in floral morphological traits across ploidy levels and specifically in corolla length across ploidy levels and genetic groups.* A, Violin plots showing the effect of sex in variation for multiple morphological traits across ploidy levels; B, corolla length differences between diploid (2x) and tetraploid (4x) individuals, shown separately for females (left) and hermaphrodites (right); C, corolla length variation among major genetic groups (Algarve, Cádiz, Donyana, Hercynian, and Tetraploid) for females (left) and hermaphrodites (right).Violin plots show kernel density, with embedded boxplots representing medians and interquartile ranges; points indicate outlier data. Asterisks in A and B indicate significance levels (ns, not significant; *, p < 0.05; ***, p < 0.001). Letters in C represent groups that are significantly different according to post-hoc comparisons on the Dunn test.

**Table S1**. Populations of Thymus sect. Mastichina, indicating their genetic cluster (Tetraploid, Hercynian, Algarve, Doñana, and Cadiz), ploidy level (estimated through flow cytometry; García-Cárdenas et al., 2025), collection site (locality, coordinates), and the herbarium voucher number. Populations are ordered alphabetically according to our field code.

| Pop. | Genetic group | Ploidy | Locality | Coordinates | Herbarium voucher (SEV) |
| --- | --- | --- | --- | --- | --- |
| A01 | Cádiz | 2x | Spain. Cádiz, Puerto Real. Dehesa de las Yeguas | 36.5557  -6.1351 | SEV290208 |
| A02 | Cádiz | 2x | Spain. Cádiz, la Barrosa Periurban Park | 36.3628  -6.1704 | SEV290209 |
| A03 | Cádiz | 2x | Spain. Cádiz, Conil de la Frontera. ZEC Pinar de Roche | 36.3163  -6.1333 | SEV290210 |
| A04 | Doñana | 2x | Spain. Sevilla. Villamanrique de la Condesa. Doñana Natural Park. La Juncosilla | 37.1916  -6.3294 | SEV290211 |
| A05 | Algarve | 2x | Portugal. Algarve, Faro. Ria Formosa Natural Park | 37.0288  -7.9773 | SEV290212 |
| A06 | Doñana | 2x | Spain. Huelva, Villarrasa. Dehesa del Duque | 37.3563  -6.6044 | SEV290213 |
| A07 | Doñana | 2x | Spain. Huelva, Bonares | 37.2586  -6.7254 | SEV290214 |
| A08 | Doñana | 2x | Spain. Doñana Natural Park. Pinar de la Algaida | 36.8553  -6.3075 | SEV290215 |
| A09 | Algarve | 2x | Portugal. Algarve, Quarteira. Vale do Lobo | 37.0617  -8.0796 | SEV290216 |
| A10 | Algarve | 2x | Portugal. Algarve, Almancil. Vale do Garrao. Ria Formosa Natural Park | 37.0443  -8.0382 | SEV290217 |
| A11 | Doñana | 2x | Spain. Huelva, Niebla. Rabo Conejo | 37,4887  -6,7192 | SEV290218 |
| A12 | Cádiz | 2x | Spain. Cádiz, Vejer de la Frontera. Las Lomas | 36.2988  -5.9087 | SEV290219 |
| A13 | Cádiz | 2x | Spain. Cádiz, Vejer de la Frontera. El Soto | 36.2422  -5.9287 | SEV290220 |
| A14 | Cádiz | 2x | Spain. Cádiz, Chiclana. Pinar del Hierro | 36.3880  -6.1222 | SEV290221 |
| A15 | Doñana | 2x | Spain. Sevilla. Villamanrique de la Condesa. Doñana Natural Park. Dehesa de Gato | 37.2540  -6.3395 | SEV290222 |
| D01 | Doñana | 2x | Spain. Huelva, Almonte. El Porretal. ZEC Doñana North and West | 37.1671  -6.4967 | SEV290223 |
| D02 | Doñana | 2x | Spain. Huelva, Hinojos. ZEC Doñana North and West | 37.2865  -6.4453 | SEV290224 |
| D04 | Algarve | 2x | Portugal. Algarve, Quelfes. Pinheiros do Marim. Ria Formosa Natural Park | 37.0336  -7.8181 | SEV290226 |
| D05 | Doñana | 2x | Spain. Huelva, Abalario. Doñana Natural Park | 37.0719  -6.6506 | SEV290227 |
| D06 | Doñana | 2x | Spain. Huelva, Ribetehilos. Doñana Natural Park | 37.1500  -6.6433 | SEV290228 |
| D07 | Doñana | 2x | Spain. Huelva, Corral de la Liebre. Doñana National Park | 36.9079  -6.4141 | SEV290229 |
| D08 | Doñana | 2x | Spain. Huelva. Corralillo Oscuro. Doñana National Park | 36.9869  -6.5215 | SEV290230 |
| M01 | Tetraploid | 4x | Spain. Sevilla, Cazalla de la Sierra. Sierra Morena de Sevilla Natural Park | 37.9075  -5.8023 | SEV290231 |
| M02 | Hercynian | 2x | Spain. Córdoba, El Vacar. ZEC Guadalmellato | 38.1001  -4.8330 | SEV290232 |
| M03 | Doñana | 2x | Spain. Huelva, Cartaya. Campo Común de Abajo | 37.2383  -7.0796 | SEV290233 |
| M04 | Tetraploid | 4x | Spain. Sevilla, Gilena | 37.2805  -4.9014 | SEV290234 |
| M05 | Tetraploid | 4x | Spain. Sevilla, Gerena | 37.5611  -6.1593 | SEV290235 |
| M06 | Tetraploid | 4x | Portugal. Algarve, Monte Gordo | 37.1855  -7.4880 | SEV290236 |
| M07 | Tetraploid | 4x | Spain. Madrid. Hoyos de Manzanares. Cuenca Alta del Manzanares Regional Park | 40.6021  -3.9167 | SEV290237 |
| M08 | Tetraploid | 4x | Spain. Cádiz, Villaluenga del Rosario. Sierra de Grazalema Natural Park | 36.8108  -5.3907 | SEV290238 |
| M09 | Tetraploid | 4x | Spain. Málaga, Ronda, Pinsapar de la Nava. Sierra de las Nieves Natural Park | 36.6650  -5.0510 | SEV290239 |
| M10 | Doñana | 2x | Spain. Huelva, Cartaya. Campo Común de Arriba | 37.3299  -7.0997 | SEV290240 |
| M11 | Doñana | 2x | Spain. Huelva, El Portil | 37.2035  -7.0136 | SEV290241 |
| M12 | Tetraploid | 4x | Spain. Cuenca, Pajaroncillo | 39.9301  -1.7257 | SEV290242 |
| M13 | Tetraploid | 4x | Spain. Cuenca, San Clemente | 39.3376  -2.4718 | SEV290243 |
| M14 | Hercynian | 2x | Portugal. Guarda, Serra da Estrela Natural Park | 40.3788  -7.5066 | SEV290244 |
| M15 | Tetraploid | 4x | Spain. León, Ponferrada | 42.5639  -6.7705 | SEV290245 |
| M16 | Tetraploid | 4x | Spain. Zamora, Toro | 41.5160  -5.4148 | SEV290246 |
| M18 | Tetraploid | 4x | Spain. Albacete, Parideras | 38.5817  -2.5262 | SEV290247 |

**Table S2.** Descriptive statistics (N, mean, standard deviation – SD, standard error – SE, and confidence intervarl – CI) for morphological traits measured individuals of *Thymus* sect. *Mastichina*. Statistics are calculated for ploidy levels and for the genetic groups identified by García-Cárdenas et al. (2025): Algarve, Cadiz, Donyana, Hercynian, and Tetraploid. All measurements are in millimeters (mm), except for pollen and stomatal size, which are given in micrometers (µm).

| Variable | Genetic group | N | min | max | mean | SD | SE | CI |
| --- | --- | --- | --- | --- | --- | --- | --- | --- |
| co | 4x | 328 | 2,28 | 7,31 | 4,27 | 1,02 | 0,06 | 0,11 |
|  | 2x | 637 | 1,76 | 6,73 | 3,52 | 0,84 | 0,03 | 0,07 |
|  | Tetraploid | 328 | 2,28 | 7,31 | 4,27 | 1,02 | 0,06 | 0,11 |
|  | Hercynian | 44 | 2,43 | 5,77 | 3,77 | 0,81 | 0,12 | 0,25 |
|  | Algarve | 92 | 2,33 | 4,87 | 3,43 | 0,56 | 0,06 | 0,12 |
|  | Doñana | 355 | 1,8 | 6,73 | 3,78 | 0,84 | 0,04 | 0,09 |
|  | Cádiz | 146 | 1,76 | 4,56 | 2,86 | 0,57 | 0,05 | 0,09 |
| ca | 4x | 328 | 1,96 | 7,08 | 4,64 | 0,84 | 0,05 | 0,09 |
|  | 2x | 637 | 1,82 | 9,05 | 3,71 | 1,07 | 0,04 | 0,08 |
|  | Tetraploid | 328 | 1,96 | 7,08 | 4,64 | 0,84 | 0,05 | 0,09 |
|  | Hercynian | 44 | 3,16 | 6,27 | 4,69 | 0,71 | 0,11 | 0,21 |
|  | Algarve | 92 | 2,16 | 4,84 | 3,14 | 0,6 | 0,06 | 0,12 |
|  | Doñana | 355 | 2,28 | 9,05 | 4,11 | 1,07 | 0,06 | 0,11 |
|  | Cádiz | 146 | 1,82 | 3,76 | 2,81 | 0,41 | 0,03 | 0,07 |
| ltooth | 4x | 328 | 1,39 | 5,03 | 3,04 | 0,62 | 0,03 | 0,07 |
|  | 2x | 637 | 0,68 | 7,08 | 2,21 | 0,85 | 0,03 | 0,07 |
|  | Tetraploid | 328 | 1,39 | 5,03 | 3,04 | 0,62 | 0,03 | 0,07 |
|  | Hercynian | 44 | 2,1 | 4,41 | 3,13 | 0,54 | 0,08 | 0,16 |
|  | Algarve | 92 | 0,68 | 2,95 | 1,73 | 0,39 | 0,04 | 0,08 |
|  | Doñana | 355 | 1,43 | 7,08 | 2,49 | 0,89 | 0,05 | 0,09 |
|  | Cádiz | 146 | 1,04 | 2,39 | 1,58 | 0,25 | 0,02 | 0,04 |
| hair | 4x | 326 | 0,22 | 1,22 | 0,71 | 0,17 | 0,01 | 0,02 |
|  | 2x | 636 | 0,11 | 1,15 | 0,41 | 0,15 | 0,01 | 0,01 |
|  | Tetraploid | 326 | 0,22 | 1,22 | 0,71 | 0,17 | 0,01 | 0,02 |
|  | Hercynian | 44 | 0,6 | 1,15 | 0,8 | 0,13 | 0,02 | 0,04 |
|  | Algarve | 91 | 0,22 | 0,54 | 0,33 | 0,07 | 0,01 | 0,01 |
|  | Doñana | 355 | 0,23 | 0,74 | 0,43 | 0,1 | 0,01 | 0,01 |
|  | Cádiz | 146 | 0,11 | 0,7 | 0,31 | 0,1 | 0,01 | 0,02 |
| fl_infl | 4x | 318 | 10 | 68 | 37,88 | 14,13 | 0,79 | 1,56 |
|  | 2x | 637 | 7 | 79 | 34,68 | 14,41 | 0,57 | 1,12 |
|  | Tetraploid | 318 | 10 | 68 | 37,88 | 14,13 | 0,79 | 1,56 |
|  | Hercynian | 44 | 25 | 79 | 41,5 | 13,49 | 2,03 | 4,1 |
|  | Algarve | 92 | 18 | 71 | 36,28 | 15,78 | 1,65 | 3,27 |
|  | Doñana | 355 | 7 | 69 | 34,16 | 14,93 | 0,79 | 1,56 |
|  | Cádiz | 146 | 9 | 61 | 32,87 | 11,68 | 0,97 | 1,91 |
| d_infl | 4x | 318 | 7,79 | 18,24 | 13,44 | 2,05 | 0,12 | 0,23 |
|  | 2x | 634 | 5,64 | 18,59 | 9,55 | 2,28 | 0,09 | 0,18 |
|  | Tetraploid | 318 | 7,79 | 18,24 | 13,44 | 2,05 | 0,12 | 0,23 |
|  | Hercynian | 44 | 9,89 | 16,29 | 11,85 | 1,37 | 0,21 | 0,42 |
|  | Algarve | 92 | 6,44 | 10,1 | 8,53 | 0,94 | 0,1 | 0,19 |
|  | Doñana | 352 | 6,22 | 18,59 | 10,31 | 2,36 | 0,13 | 0,25 |
|  | Cádiz | 146 | 5,64 | 10,57 | 7,68 | 0,94 | 0,08 | 0,15 |
| prop_ltooth | 4x | 328 | 0,48 | 0,84 | 0,65 | 0,06 | 0 | 0,01 |
|  | 2x | 637 | 0,26 | 0,81 | 0,59 | 0,06 | 0 | 0,01 |
|  | Tetraploid | 328 | 0,48 | 0,84 | 0,65 | 0,06 | 0 | 0,01 |
|  | Hercynian | 44 | 0,53 | 0,75 | 0,67 | 0,04 | 0,01 | 0,01 |
|  | Algarve | 92 | 0,26 | 0,74 | 0,55 | 0,07 | 0,01 | 0,01 |
|  | Doñana | 355 | 0,47 | 0,81 | 0,6 | 0,06 | 0 | 0,01 |
|  | Cádiz | 146 | 0,46 | 0,72 | 0,56 | 0,04 | 0 | 0,01 |
| d_max | 4x | 70 | 21 | 60 | 34,66 | 7,34 | 0,88 | 1,75 |
|  | 2x | 30 | 25 | 42 | 35,93 | 3,33 | 0,61 | 1,24 |
|  | Tetraploid | 70 | 21 | 60 | 34,66 | 7,34 | 0,88 | 1,75 |
|  | Hercynian | 20 | 25 | 42 | 36,2 | 3,62 | 0,81 | 1,69 |
|  | Doñana | 10 | 30 | 39 | 35,4 | 2,76 | 0,87 | 1,97 |
| d_min | 4x | 70 | 12 | 51 | 29,61 | 8,03 | 0,96 | 1,91 |
|  | 2x | 30 | 19 | 40 | 30,83 | 4,82 | 0,88 | 1,8 |
|  | Tetraploid | 70 | 12 | 51 | 29,61 | 8,03 | 0,96 | 1,91 |
|  | Hercynian | 20 | 19 | 40 | 31,25 | 5,39 | 1,2 | 2,52 |
|  | Doñana | 10 | 24 | 37 | 30 | 3,53 | 1,12 | 2,52 |
| stoma_w | 4x | 30 | 10 | 21 | 15,33 | 2,43 | 0,44 | 0,91 |
|  | 2x | 180 | 9 | 25 | 14,89 | 2,61 | 0,19 | 0,38 |
|  | Tetraploid | 30 | 10 | 21 | 15,33 | 2,43 | 0,44 | 0,91 |
|  | Hercynian | 60 | 13 | 20 | 15,93 | 1,88 | 0,24 | 0,48 |
|  | Algarve | 30 | 10 | 22 | 14,5 | 2,45 | 0,45 | 0,91 |
|  | Doñana | 60 | 9 | 25 | 15,07 | 2,83 | 0,37 | 0,73 |
|  | Cádiz | 30 | 9 | 18 | 12,83 | 2,38 | 0,43 | 0,89 |
| stoma_l | 4x | 30 | 15 | 32 | 22,23 | 4,53 | 0,83 | 1,69 |
|  | 2x | 180 | 11 | 32 | 20,02 | 3,96 | 0,3 | 0,58 |
|  | Tetraploid | 30 | 15 | 32 | 22,23 | 4,53 | 0,83 | 1,69 |
|  | Hercynian | 60 | 11 | 32 | 21,85 | 4,68 | 0,6 | 1,21 |
|  | Algarve | 30 | 14 | 25 | 18,87 | 2,37 | 0,43 | 0,89 |
|  | Doñana | 60 | 11 | 30 | 19,97 | 3,56 | 0,46 | 0,92 |
|  | Cádiz | 30 | 12 | 24 | 17,63 | 2,57 | 0,47 | 0,96 |

**See attached excel file.**

**Table S3.** Descriptive statistics [sample size (N), mean, standard deviation (SD), standard error (SE), and confidence interval (CI)] for morphological traits measured individuals of *Thymus* sect. *Mastichina*. Statistics are calculated for all sampled populations. All measurements are in millimeters (mm), except for pollen and stomatal size, which are given in micrometers (µm).
